# Elicitation of broadly protective immunity to influenza by multivalent hemagglutinin nanoparticle vaccines

**DOI:** 10.1101/2020.05.30.125179

**Authors:** Seyhan Boyoglu-Barnum, Daniel Ellis, Rebecca A. Gillespie, Geoffrey B. Hutchinson, Young-Jun Park, Syed M. Moin, Oliver Acton, Rashmi Ravichandran, Mike Murphy, Deleah Pettie, Nick Matheson, Lauren Carter, Adrian Creanga, Michael J. Watson, Sally Kephart, John R. Vaile, George Ueda, Michelle C. Crank, Lance Stewart, Kelly K. Lee, Miklos Guttman, David Baker, John R. Mascola, David Veesler, Barney S. Graham, Neil P. King, Masaru Kanekiyo

## Abstract

Influenza vaccines that confer broad and durable protection against diverse virus strains would have a major impact on global health. However, next-generation vaccine design efforts have been complicated by challenges including the genetic plasticity of the virus and the immunodominance of certain epitopes in its glycoprotein antigens. Here we show that computationally designed, two-component nanoparticle immunogens induce potently neutralizing and broadly protective antibody responses against a wide variety of influenza viruses. The nanoparticle immunogens display 20 hemagglutinin (HA) trimers in a highly immunogenic array, and their assembly *in vitro* enables precisely controlled co-display of multiple distinct HAs in defined ratios. Nanoparticle immunogens displaying the four HAs of licensed quadrivalent influenza vaccines (QIV) elicited hemagglutination inhibition and neutralizing antibody responses to vaccine-matched strains that were equivalent or superior to commercial QIV in mice, ferrets, and nonhuman primates. The nanoparticle immunogens—but not QIV—simultaneously induced broadly protective antibody responses to heterologous viruses, including H5N1 and H7N9, by targeting the subdominant yet conserved HA stem. Unlike previously reported influenza vaccine candidates, our nanoparticle immunogens can alter the intrinsic immunodominance hierarchy of HA to induce both potent receptor-blocking and broadly cross-reactive stem-directed antibody responses and are attractive candidates for a next-generation influenza vaccine that could replace current seasonal vaccines.

**One Sentence Summary:** Nanoparticle immunogens displaying four seasonal influenza hemagglutinins elicit neutralizing antibodies directed at both the immunodominant head and the conserved stem and confer broad protective immunity.

## Introduction

Influenza viruses cause an estimated 290,000–650,000 deaths annually despite the availability of licensed vaccines^1^. Strains currently circulating in humans belong to the H1N1 and H3N2 subtypes of group 1 and 2 influenza A viruses, respectively, as well as the Yamagata (B/Yam) and Victoria (B/Vic) lineages of influenza B^2,3^. Most current seasonal influenza vaccines comprise antigens each of these four branches and provide protection against symptomatic influenza infection ranging from about 60% down to less than 10%, varying from year to year^4^. Vaccine efficacy largely derives from eliciting antibodies to hemagglutinin (HA), the immunodominant surface glycoprotein of influenza viruses. Current vaccines induce narrow, strain-specific responses predominantly targeting the hypervariable head domain of HA, and lose efficacy from one season to the next due to antigenic variation. Vaccine performance is also blunted when antigenic mismatches occur between the vaccine and circulating strains arising from mispredictions or egg-adapted mutations introduced during vaccine manufacturing^5–9^. These challenges require continual updating of the vaccine strains and annual vaccine reformulation. Furthermore, current seasonal vaccines do not protect against pandemic strains or subtypes such as H5N1 or H7N9^10–12^. Four influenza pandemics have occurred since 1900 with varying levels of severity^13^, including the 1918 Spanish flu which claimed 50–100 million lives^14^ and the recent 2009 H1N1 pandemic. In the absence of novel interventions, it is inevitable that they will continue to reoccur^15–17^.

HA is a homotrimeric class I fusion protein, consisting of a head domain that facilitates host cell attachment by binding cell-surface sialosides and a stem domain containing the membrane fusion machinery^18,19^. Both the head and stem contain highly conserved epitopes that are targeted by broadly neutralizing antibodies (bnAbs) capable of protecting against infection by diverse influenza viruses in animal models^18,20,21^. A few such bnAbs are currently being evaluated in human clinical trials for their prophylactic and therapeutic potential^22^. However, bnAbs are rarely elicited by immunization or infection. Instead, antibodies targeting immunodominant hypervariable epitopes in the head domain often dominate the response to HA, resulting in strain-specific immunity and limited breadth of protection^23–25^. Because of the subdominant nature and broader cross-reactive potential of epitopes in the HA stem, several antigen design efforts have focused on removing or obscuring the head domain to elicit improved stem-focused responses^26–32^.

Orderly arrays of antigen on submicron particles are efficiently recognized by the immune system and induce robust humoral responses^33^. The utility of non-viral self-assembling proteins for displaying complex antigens was initially demonstrated by displaying native-like HA trimers on *H. pylori* ferritin nanoparticles, which increased the overall immunogenicity of HA and improved the breadth of vaccine-elicited antibody responses^34^. To further improve breadth across multiple homotypic and heterosubtypic strains, ferritin-based display has been combined with re-designed HA antigens, including headless trimeric HA stems and antigenically optimized HAs^26,28,29,35^. More recently, ferritin was used as a scaffold for co-displaying the monomeric receptor-binding domains (RBDs) from multiple H1 HAs on the same nanoparticle surface^36^. These “mosaic” nanoparticles elicited antibody responses of greater breadth and potency than corresponding mixtures or “cocktails” of nanoparticles each displaying individual RBDs. However, spontaneous assembly of the homomeric ferritin scaffold in cells during secretion precludes the production of mosaic nanoparticles co-displaying multiple trimeric HA antigens.

Recent advances in computational design methods have made possible the predictive design of novel self-assembling protein complexes with structures tailored to specific applications^37–42^. In addition to designed homomeric assemblies of exceptional stability, designed two-component complexes constructed from multiple copies of two distinct protein building blocks have been used to generate nanoparticle immunogens. Multivalent antigen display on these nanoparticles enhanced the potency of vaccine-elicited immune responses against a malaria antigen^43^ as well as two trimeric class I viral fusion proteins: prefusion-stabilized respiratory syncytial virus F and HIV envelope^44–46^. Assembly of the two-component nanoparticles *in vitro* from independently purified components could in theory enable facile, scalable, and stoichiometrically-controlled co-display of multiple antigenic variants of a wide variety of oligomeric antigens.

Here we use *in vitro* assembly to produce cocktail and mosaic nanoparticle immunogens (co-)displaying multiple trimeric HA ectodomains and evaluate their immunogenicity and protective efficacy against diverse influenza viruses in mice, ferrets, and nonhuman primates. We find that these immunogens possess unique antigenic and immunogenic properties, and have the potential to induce protective responses against seasonal and pandemic strains effective over multiple years without reformulation.

### Immunogen design, *in vitro* assembly, and characterization

We genetically fused HA ectodomains from the four strains in licensed 2017-2018 seasonal influenza vaccines to the N terminus of I53_dn5B, the trimeric component of the two-component icosahedral nanoparticle I53_dn5 (Fig. 1a and Supplementary Table 1; H1: A/Michigan/45/2015, H3: A/Hong Kong/4801/2014, B/Yam: B/Phuket/3073/2013, B/Vic: B/Brisbane/60/2008; ref. 42). The four proteins were individually produced in mammalian cells and purified by affinity chromatography and size-exclusion chromatography (SEC). The SEC profiles were consistent with the expected size of the trimeric fusion proteins (Extended Data Fig. 1a), and SDS-PAGE indicated that the preparations contained exclusively uncleaved HA-I53_dn5B fusion proteins (Extended Data Fig. 1b and Supplementary Fig. 1a). The antigenic structure of each HA was confirmed by biolayer interferometry (BLI) using monoclonal antibodies (mAbs) with appropriate specificities (Extended Data Fig. 1c).

**Figure 1.**
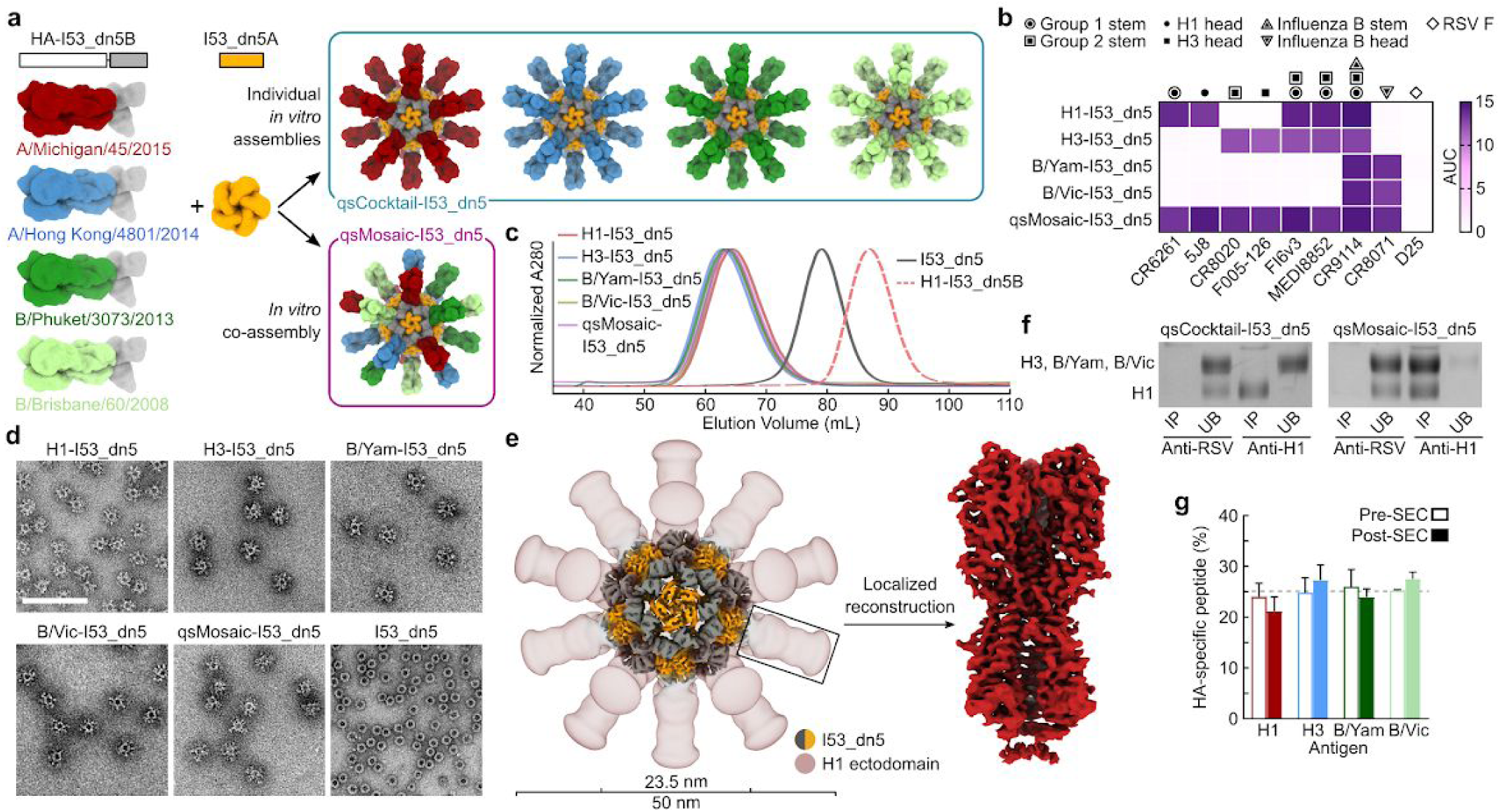
Design, *in vitro* assembly, and characterization of cocktail and mosaic HA nanoparticle immunogens. **a**, Schematics of the components and *in vitro* assembly of nanoparticle immunogens displaying HA. Trimeric HA-I53_dn5B fusions were secreted from HEK293F cells, while the I53_dn5A pentamer was expressed in *E. coli*. **b**, Antigenic characterization of purified nanoparticle immunogens by ELISA. Symbols indicate the specificity of each mAb. AUC, area under the curve. **c**, Analytical SEC of purified nanoparticle immunogens, compared to I53_dn5 nanoparticles lacking displayed antigen and trimeric H1-I53_dn5B, using a Sephacryl S-500 HR 16/60 column. **d**, Negative stain electron micrographs of purified nanoparticle immunogens and I53_dn5 (scale bar, 200 nm). **e**, 3D reconstruction of the H1-I53_dn5 nanoparticle immunogen and local reconstruction of H1 MI15 obtained using single-particle cryo-EM. For the nanoparticle immunogen, high (grey/orange) and low (red) contour 3D representations are overlaid to enable visualization of the I53_dn5 scaffold and the displayed HA, respectively. The density corresponding to the low contour representation was smoothed using a 16 Å low-pass filter for clarity. **f**, Immunoprecipitation of qsCocktail-I53_dn5 and qsMosaic-I53_dn5 using RSV F-specific (MPE8) and H1-specific (5J8) mAbs. IP, immunoprecipitated; UB, unbound. **g**, Quantitation of HA antigen content in assembled qsMosaic-I53_dn5 nanoparticles before and after preparative SEC by peptide mass spectrometry. The dashed grey line represents the expected peptide content from each HA, and error bars represent the s.d. of measurements across four unique peptides from each HA. The peptides used to quantify each HA are provided in Supplementary Table 3.

To produce a mosaic nanoparticle co-displaying quadrivalent seasonal HAs (qsMosaic-I53_dn5), the purified trimeric HA-I53_dn5B components were mixed in equimolar amounts prior to addition of purified I53_dn5A pentamer to induce nanoparticle assembly (Fig. 1a). In parallel, nanoparticles individually displaying each HA ectodomain were produced by separately assembling each of the four HA-I53_dn5B trimers with I53_dn5A pentamer. During the purification of each nanoparticle by SEC, nearly all of the protein eluted in an early peak corresponding to the assembled icosahedral complex with only minor amounts of residual, unassembled components (Extended Data Fig. 1d,e and Supplementary Fig. 1b). The nanoparticles bound head- and stem-directed mAbs specific to the HA trimers displayed, including qsMosaic-I53_dn5, which bound mAbs specific to H1, H3, and influenza B HAs (Fig. 1b). The size and morphology of each nanoparticle, including I53_dn5 nanoparticles without HA, was evaluated by analytical SEC (Fig. 1c), dynamic light scattering (Extended Data Fig. 1f), and negative stain electron microscopy (Fig. 1d), which confirmed assembly to the intended icosahedral architecture with no evidence of aggregation. A single-particle cryo-EM reconstruction of the H1-I53_dn5 nanoparticle to 6.6 Å resolution established the integrity of the I53_dn5 nanoparticle core and indicated marked flexibility between the displayed HA ectodomains and the underlying nanoparticle scaffold despite the use of a minimal (two-residue) linker (Fig. 1e and Extended Data Fig. 1g-i). We therefore used localized reconstruction^47^ to determine a 3.3 Å reconstruction of the displayed H1 HA, demonstrating full retention of the native structure of the genetically fused antigen (Fig. 1e and Extended Data Fig. 1g,j). Finally, hydrogen-deuterium exchange mass spectrometry revealed no significant differences in the local structural order of H1-I53_dn5 and an H1-foldon fusion protein, a well-characterized research reagent^48^, further confirming at high resolution the integrity of the HA displayed on the nanoparticle (Extended Data Fig. 2). These data demonstrate that *in vitro* assembly yielded monodisperse I53_dn5-based nanoparticles with the expected size, morphology, and antigenicity.

We prepared a quadrivalent seasonal cocktail containing equimolar amounts of the four individual HA-displaying nanoparticles (qsCocktail-I53_dn5; Fig. 1a) and used two distinct assays to confirm co-assembly in the case of qsMosaic-I53_dn5. First, we immunoprecipitated each nanoparticle using the H1-specific mAb 5J8 (ref. 49) and exploited the unique mobility of the H1-I53_dn5B band during non-reducing SDS-PAGE to track the fate of H1-containing nanoparticles. While immunoprecipitation of qsCocktail-I53_dn5 nanoparticles recovered only H1-I53_dn5B and not the other HA-I53_dn5B proteins, the same procedure resulted in complete pull-down of all the HA-I53_dn5B proteins in qsMosaic-I53_dn5 nanoparticles (Fig. 1f and Supplementary Fig. 1c). No pull-down was seen when immunoprecipitation was performed with an irrelevant, RSV F-specific mAb. This result indicates that co-assembly of qsMosaic-I53_dn5 is very efficient—as expected if there is no preferential incorporation or exclusion of specific HAs during assembly (Extended Data Fig. 3a,b)—and confirms that subunit exchange did not occur in the qsCocktail-I53_dn5 nanoparticle. Second, we devised a sandwich BLI experiment in which each nanoparticle was captured by immobilized H1-specific mAb and subsequently evaluated for binding to H3- or influenza B-specific mAbs. qsMosaic-I53_dn5 nanoparticles were bound by the H3- and B-specific mAbs, indicating co-display, while qsCocktail-I53_dn5 nanoparticles were not (Extended Data Fig. 3c).

We used quantitative mass spectrometry to measure the relative amount of each HA in co-assembled qsMosaic-I53_dn5 nanoparticles before and after purification by SEC. Quantitating unique peptides for each HA antigen revealed that the equimolar HA content used in the *in vitro* assembly reaction was maintained in the purified nanoparticle (Fig. 1g and Supplementary Table 3). Post-SEC specific HA content was also maintained in additional mosaic nanoparticles in which we substantially altered the stoichiometric ratios of the four HA-I53_dn5B components (Extended Data Fig. 3d). Together, these data demonstrate that *in vitro* assembly enables co-display of multiple oligomeric antigens in defined ratios. We note, however, that the symmetric nature of the I53_dn5 nanoparticle scaffold does not allow the controlled placement of antigens in specific locations on the nanoparticle, and we expect that the antigen content of each individual nanoparticle will assume a distribution centered on the overall antigen content (Extended Data Fig. 3a).

### Vaccine-elicited antibody responses against vaccine-matched antigens and viruses

We next performed a series of *in vivo* experiments in mice, ferrets, and nonhuman primates (NHPs) to compare the immunogenicity of qsCocktail-I53_dn5 and qsMosaic-I53_dn5 to commercial 2017-2018 QIV (Fig. 2a). Throughout our studies, we matched the total protein dose of each nanoparticle immunogen to the HA content of QIV. After three immunizations with each immunogen formulated with AddaVax, a squalene-based oil-in-water emulsion chemically equivalent to the licensed adjuvant MF59 (ref. 50), we measured antigen-specific antibody, hemagglutination inhibition (HAI), and microneutralization titers against vaccine-matched or slightly mismatched viruses and HAs (Fig. 2b). HA-specific antibody titers induced by both nanoparticle immunogens were equivalent or superior to those induced by QIV, with larger increases observed in mice and ferrets than in NHPs in most cases (Extended Data Fig. 4). Similarly, in mice and ferrets qsCocktail-I53_dn5 and qsMosaic-I53_dn5 elicited higher HAI titers than QIV against vaccine-matched B/Yam virus and were comparable against H1N1 and a slightly mismatched B/Vic virus (Fig. 2c). The three immunogens induced roughly equivalent HAI activity against these viruses in NHPs. The increases in microneutralization titer afforded by the nanoparticle immunogens over QIV were more variable, in some cases exceeding an order of magnitude, but overall exhibited the same trend of being comparable to each other and equivalent or superior to QIV (Fig. 2d). Additional immunogenicity studies in mice without adjuvant (Extended Data Fig. 5) and using updated versions of the three immunogens containing the 2018-2019 vaccine strains (Extended Data Fig. 6a-d) yielded similar results. Together, these data establish that qsCocktail-I53_dn5 and qsMosaic-I53_dn5 elicit vaccine-matched antibody responses that are equivalent or superior to current commercial influenza vaccines.

**Figure 2.**
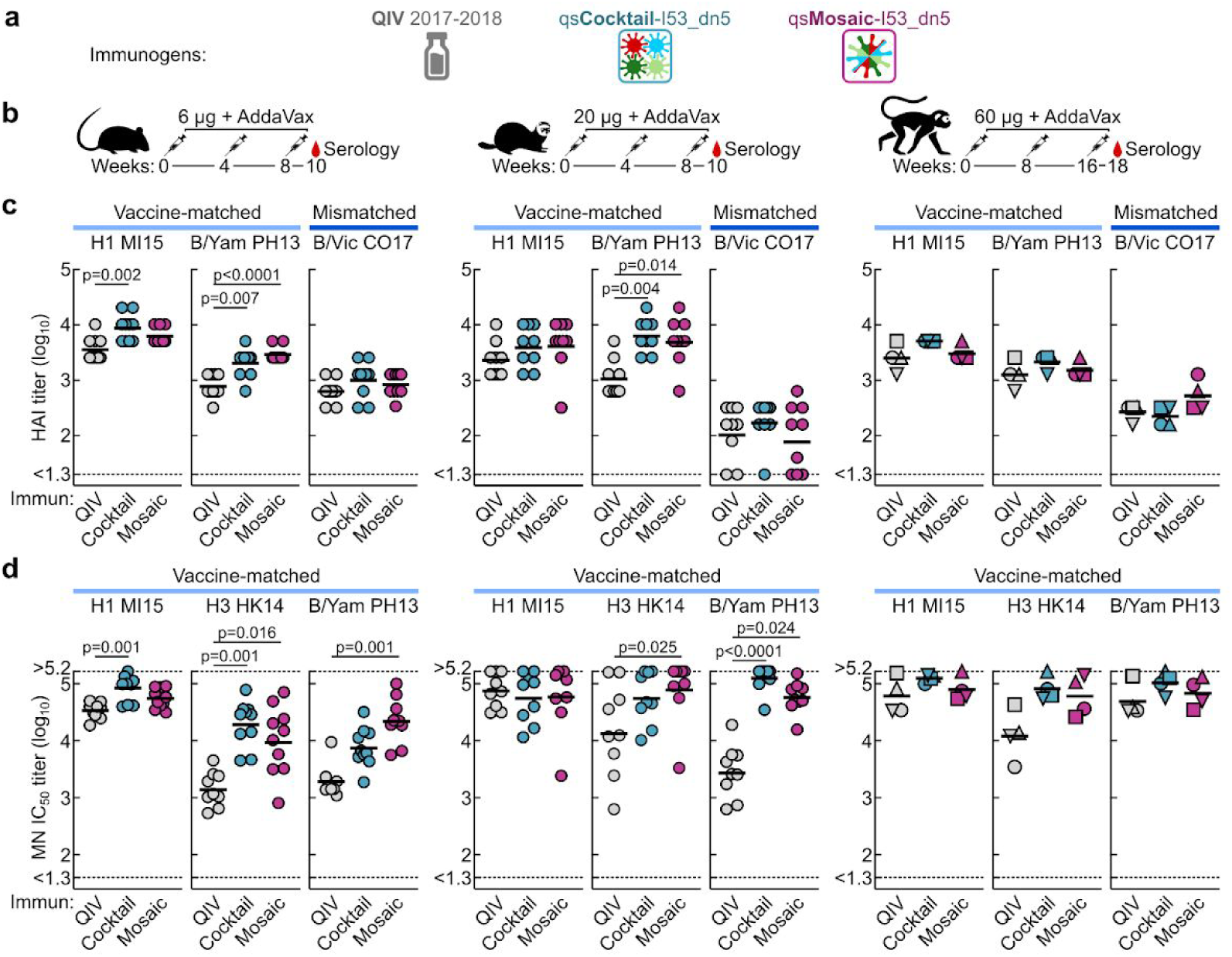
Vaccine-elicited antibody responses against vaccine-matched viruses in mice, ferrets, and NHPs. **a**, Immunogens used in the animal studies. **b**, Immunization schemes for mouse, ferret, and NHP studies. All immunizations were given intramuscularly with AddaVax. Groups of BALB/cJ mice (*N* = 10), Finch ferrets (*N* = 9), and rhesus macaques (*N* = 4) were used in each experiment. **c**, Hemagglutination inhibition (HAI) and **d**, microneutralization (MN) titers in immune sera. Microneutralization titers are reported as half maximal inhibitory dilution (IC_50_). H3N2 virus did not efficiently agglutinate red blood cells, so HAI was not measured for this virus. Each symbol represents an individual animal and the horizontal bar indicates the geometric mean of the group. Individual NHPs are identified by unique symbols. Statistical analysis was performed using nonparametric Kruskal–Wallis test with Dunn’s multiple comparisons. All animal experiments except for NHPs were performed at least twice and representative data are shown.

### Neutralization of and protection against historical H1N1 and H3N2 viruses

We next tested sera from ferrets immunized with QIV, qsCocktail-I53_dn5, and qsMosaic-I53_dn5 for their ability to neutralize a panel of H1N1 and H3N2 viruses that represent historical antigenic drift and shift^51^ (Fig. 3a and Supplementary Table 2). H1N1 microneutralization showed a clear demarcation between pre- and post-2009 viruses, with much lower neutralization activity overall against pre-2009 viruses possessing HAs antigenically distant from the vaccine-strain H1 (Fig. 3c). Nevertheless, both nanoparticle immunogens elicited roughly equivalent or superior neutralizing activity to QIV for all H1N1 strains tested, which translated to slightly higher global geometric mean titers (GMT) across 10 viruses (Fig. 3d). Historical H3N2 microneutralization decreased more gradually, in keeping with the lineage’s continuous antigenic drift, and both nanoparticle immunogens elicited ∼10-fold higher levels of neutralizing activity than QIV against viruses dating back to 2002 (Fig. 3c), resulting in significant increases in global GMT (Fig. 3e).

**Figure 3.**
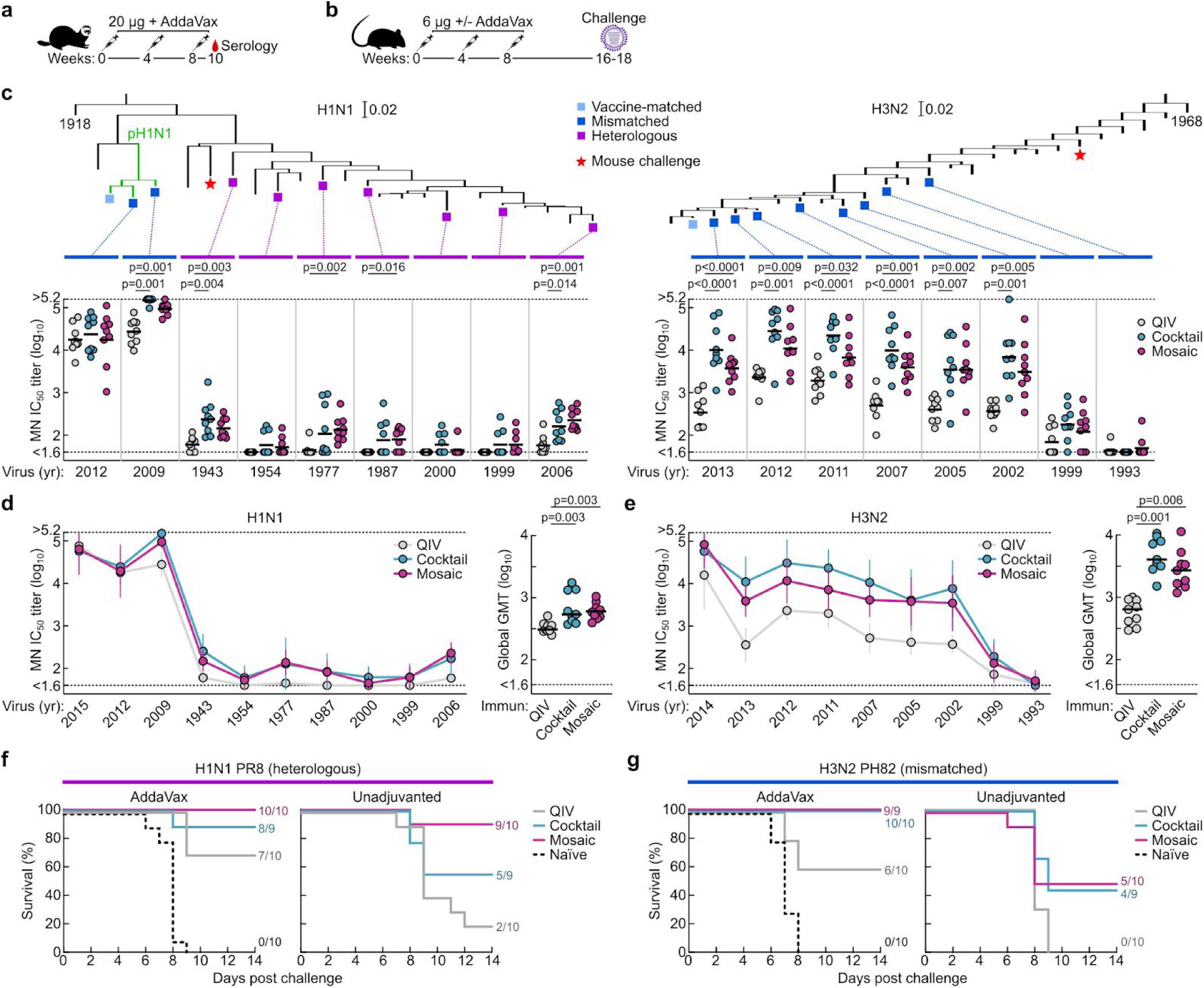
Neutralization of and protection against historical H1N1 and H3N2 viruses. **a**, Immunization scheme for ferret study. Groups of Finch ferrets (*N* = 9) were used. **b**, Immunization and virus challenge scheme for mouse studies. Groups of BALB/cJ mice (*N* = 9-10) were used in each experiment. **c**, Neutralization of representative panels of historical H1N1 and H3N2 viruses by ferret immune sera. Phylogenetic trees of HA sequences of human H1N1 (left) and H3N2 (right) viruses are shown. Viruses used in neutralization assays and mouse challenge experiments are indicated as colored squares and stars, respectively. **d–e**, Summary of neutralization breadth of ferret immune sera across multiple H1N1 and H3N2 viruses. Geometric mean IC_50_ titers ± geometric s.d. for each group are shown (left). Means of geometric mean IC_50_ titers across 10 H1N1 or 9 H3N2 viruses for each individual animal are plotted as global GMT (right). Each symbol represents an individual animal and the horizontal bar indicates the geometric mean (**c** and **d**) or mean (**d** and **e**) of the group. Statistical analysis was performed using nonparametric Kruskal–Wallis test with Dunn’s multiple comparisons. **f**, Heterologous H1N1 and **g**, mismatched H3N2 virus challenge in immunized mice. Viruses were given intranasally on day 0. Mice were monitored for 14 days post infection. Ferret experiments were performed twice and representative data are shown, while mouse challenge experiments were performed once.

Encouraged by these results, we compared the ability of QIV, qsCocktail-I53_dn5, and qsMosaic-I53_dn5 to protect against lethal challenges with highly divergent H1N1 and H3N2 viruses. Mice were immunized with AddaVax-adjuvanted or unadjuvanted immunogens and subsequently challenged with heterologous A/Puerto Rico/8/1934 (H1N1) or mismatched A/Philippines/2/1982 (H3N2) viruses (Fig. 3b,c). All mice receiving mock immunizations succumbed to disease and had to be euthanized by 9 days post-infection. When adjuvanted, both nanoparticle immunogens provided complete or near-complete protection (97% in aggregate across the four groups), while QIV afforded partial protection against both H1N1 (70%) and H3N2 (60%) challenges (Fig. 3f,g). Remarkably, in the absence of adjuvant, mice immunized with qsMosaic-I53_dn5 were almost completely protected from heterologous H1N1 challenge (90%) and partially protected from mismatched H3N2 challenge (50%), while qsCocktail-I53_dn5 provided partial protection in both cases (55% and 44%, respectively). In contrast, unadjuvanted QIV provided negligible protection (20% and 0%, respectively).

Together, these data establish that both qsCocktail-I53_dn5 and qsMosaic-I53_dn5 elicit more broadly neutralizing antibody responses and protective immunity against mismatched H1N1 and H3N2 viruses than current commercial influenza vaccines. This effect was magnified in the absence of adjuvant. These findings suggest the nanoparticle immunogens might confer multi-season protection without requiring annual vaccine reformulation.

### Vaccine-elicited heterosubtypic antibody responses and protective immunity

Current commercial influenza vaccines are limited in their ability to protect against many widely circulating zoonotic strains, making influenza a significant pandemic threat^52^. We compared the ability of QIV, qsCocktail-I53_dn5, and qsMosaic-I53_dn5 to provide immunity against heterosubtypic influenza A viruses in mice, ferrets, and NHPs. We found that both nanoparticle immunogens elicited cross-reactive antibody responses to HAs from heterosubtypic group 1 (H5N1 and H6N1) and group 2 (H7N9 and H10N8) viruses, whereas QIV elicited low—in some cases undetectable—levels of such antibodies (Fig. 4a-c). The ability of the nanoparticle immunogens to induce heterosubtypic antibody responses is not limited to the 2017-2018 vaccine composition, as we observed similar cross-reactive antibody responses in mice immunized with nanoparticle immunogens updated with the 2018-2019 vaccine strains (Extended Data Fig. 6e).

**Figure 4.**
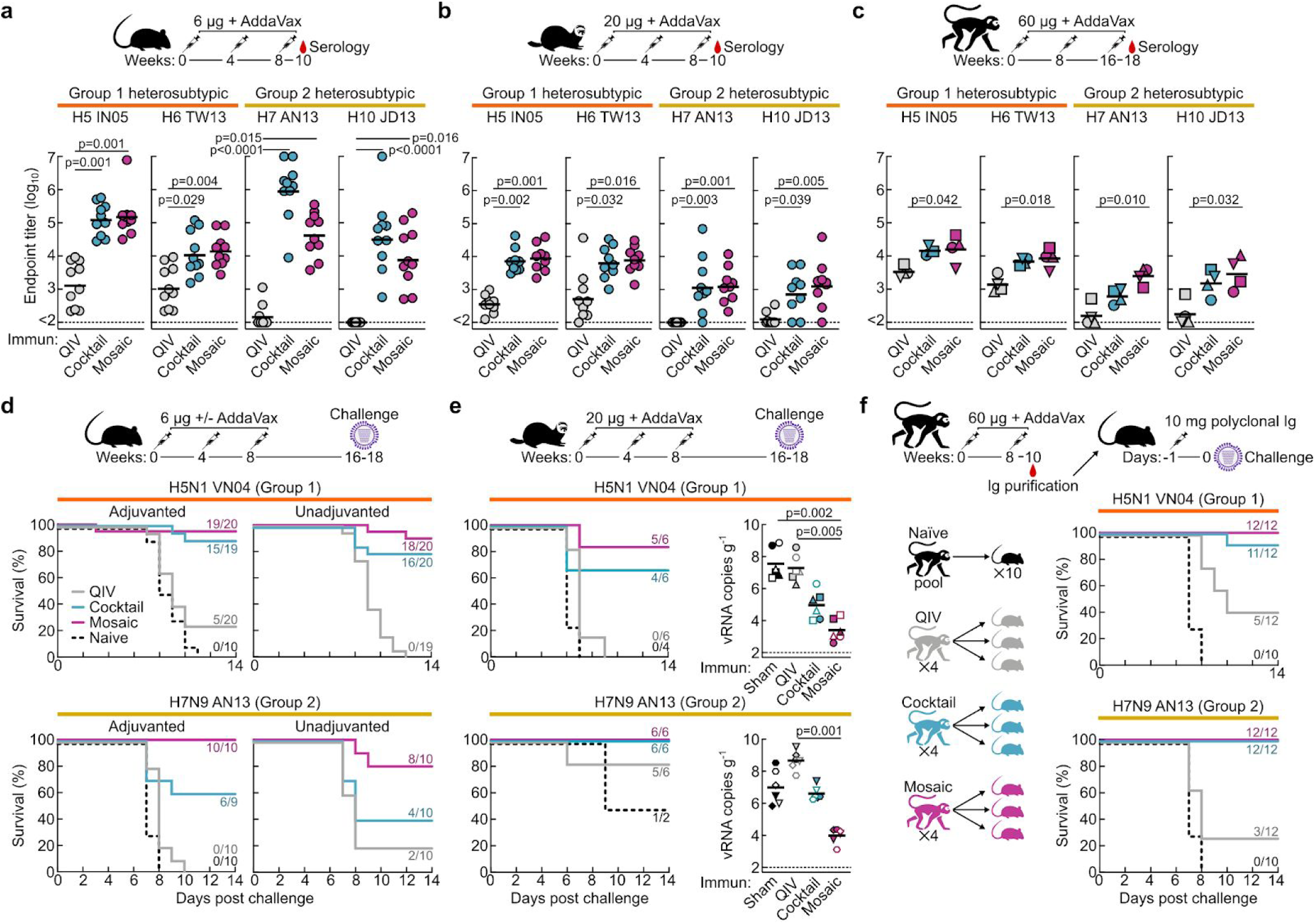
Vaccine-elicited heterosubtypic antibody responses and protective immunity. Cross-reactive antibody responses to heterosubtypic HA antigens in **a**, mice, **b**, ferrets, and **c**, NHPs. Immunization schemes are shown at the top of each panel. Antibody titers are expressed as endpoint dilutions. Each symbol represents an individual animal and the horizontal bar indicates the geometric mean of each group of BALB/cJ mice (*N* = 10), Finch ferrets (*N* = 9), and rhesus macaques (*N* = 4). Individual NHPs are identified by unique symbols. All animal immunization experiments except for NHP were performed at least twice and representative data are shown. **d**, Heterosubtypic influenza virus challenge in immunized mice. Immunized mice were challenged with either H5N1 (top) or H7N9 virus (bottom) intranasally. **e**, Heterosubtypic influenza virus challenge in immunized ferrets. Immunized ferrets were challenged with either H5N1 (top) or H7N9 virus (bottom) intranasally. Three ferrets from each group were euthanized at day 4 post challenge to measure viral RNA in lung tissue (right). Individual ferrets are identified by unique symbols. Right and left caudal lung lobes are indicated as closed and open symbols, respectively. **f**, Heterosubtypic influenza virus challenge after passive transfer of purified NHP immune Ig in mice. Polyclonal Ig from immunized NHPs was purified and administered intraperitoneally to recipient mice prior to infection with H5N1 (top) or H7N9 (bottom) virus. Control Ig was purified from pooled naïve sera and given to 10 mice. Statistical analysis was performed using nonparametric Kruskal–Wallis test with Dunn’s multiple comparisons. Mouse challenge experiments were performed twice, while ferret experiments and passive transfer experiments were performed once, and representative data are shown.

To assess whether these cross-reactive antibody responses were protective, we carried out several challenge studies with H5N1 and H7N9 viruses in mice and ferrets. First, we immunized mice with each of the three immunogens either with or without AddaVax and challenged them with lethal doses of heterosubtypic A/Vietnam/1203/2004 (H5N1) or A/Anhui/1/2013 (H7N9) virus 8-10 weeks after the last immunization. As expected, all animals receiving mock immunizations succumbed to disease, and QIV provided negligible protection (12% in aggregate across the four experiments; Fig. 4d). Strikingly, qsCocktail-I53_dn5 conferred partial protection (71% in aggregate) and qsMosaic-I53_dn5 nearly complete protection (92% in aggregate) against these heterosubtypic challenges, even in the absence of adjuvant (67% and 87%, respectively). Similar results were obtained with adjuvant in ferrets, where the nanoparticle immunogens provided robust protection against both H5N1 and H7N9 challenge (88% combined), whereas commercial QIV (33% combined) and mock immunization (17%) provided only weak protection (Fig. 4e). Viral RNA quantification in lung tissues by RT-qPCR revealed that the animals receiving qsMosaic-I53_dn5 had significantly lower H5N1 or H7N9 virus copy numbers in the lung than those immunized with commercial QIV (Fig. 4e, right).

To determine whether vaccine-elicited serum antibodies alone could confer protection against heterosubtypic challenge, we performed passive transfer experiments in mice. We passively immunized three mice with 10 mg of purified immunoglobulin (Ig) from each immunized NHP 24 h prior to infection with H5N1 or H7N9 virus (Fig. 4f). We included ten mice passively immunized with Ig purified from influenza-naïve NHPs as negative controls and another ten mice passively immunized with FI6v3, a potent human stem-directed bnAb^53^, as positive controls. All mice that received Ig from qsCocktail-I53_dn5- or qsMosaic-I53_dn5-immunized animals, as well as the animals that received FI6v3, showed no weight loss and were protected from disease, with the exception of one mouse receiving Ig from an NHP immunized with qsCocktail-I53_dn5. In contrast, all mice that received naïve Ig succumbed to disease and had to be euthanized by 8 days post-infection, while the mice that received Ig from QIV-immunized NHPs showed significant weight loss and only partial protection against H5N1 (42%) and H7N9 (25%) challenge.

To better understand the role of multivalent antigen display in the remarkable antibody responses elicited by qsCocktail-I53_dn5 and qsMosaic-I53_dn5 in mice, ferrets, and NHPs, we performed another immunization experiment in mice in which we included a non-assembling version of these immunogens. This non-assembling control immunogen was identical to qsCocktail-I53_dn5 and qsMosaic-I53_dn5 except that the trimeric components lacked the computationally designed interface that drives nanoparticle assembly (Extended Data Fig. 7a-d and Supplementary Table 1; ref. 42). While the non-assembling immunogen elicited microneutralization titers against vaccine-matched viruses that were similar to qsCocktail-I53_dn5 and qsMosaic-I53_dn5, the cross-reactive antibody responses elicited against H5N1 and H7N9 HAs were more than 10- and 100-fold lower, respectively, and were similar to those induced by commercial QIV (Extended Data Fig. 7e-g).

Together, these data demonstrate that qsCocktail-I53_dn5 and qsMosaic-I53_dn5 elicit broadly protective antibody responses against a wide variety of influenza viruses, including known pandemic threats, in contrast to currently available influenza vaccines. Formation of the nanoparticle architecture displaying an orderly array of HA antigens is required, since an equivalent dose of non-assembling HA trimeric components and the I53_dn5A pentamer elicited a negligible cross-reactive response.

### Molecular basis for vaccine-elicited broad antibody responses against heterosubtypic influenza viruses

We performed several experiments to better understand the molecular and structural determinants of the broadly protective immunity induced by qsCocktail-I53_dn5 and qsMosaic-I53_dn5. First, we measured serum antibody binding to stabilized stem trimers of group 1 and group 2 HAs, which comprise only the conserved stem domain^26,28^. Stem-directed antibodies elicited by both qsCocktail-I53_dn5 and qsMosaic-I53_dn5 were significantly higher than those induced by QIV in all three animal species (Fig. 5a). We next assessed whether these stem-directed antibodies contribute to virus neutralization by performing microneutralization assays in the presence of competitor proteins to deplete certain antibody specificities. Analysis of sera from NHPs immunized with qsMosaic-I53_dn5 revealed that neutralizing activity against the vaccine-matched H1N1 virus was depleted by vaccine-matched full-length HA ectodomain but not by an H1 HA stem, suggesting that antibodies targeting epitopes outside of the stem domain account for most of the neutralizing activity, as expected (Fig. 5b). In contrast, neutralizing activity against a heterosubtypic H5N1 virus was fully depleted by both vaccine-matched H1 HA and an H1 HA stem, indicating that stem-directed antibodies are solely responsible for the observed heterosubtypic neutralization.

**Figure 5.**
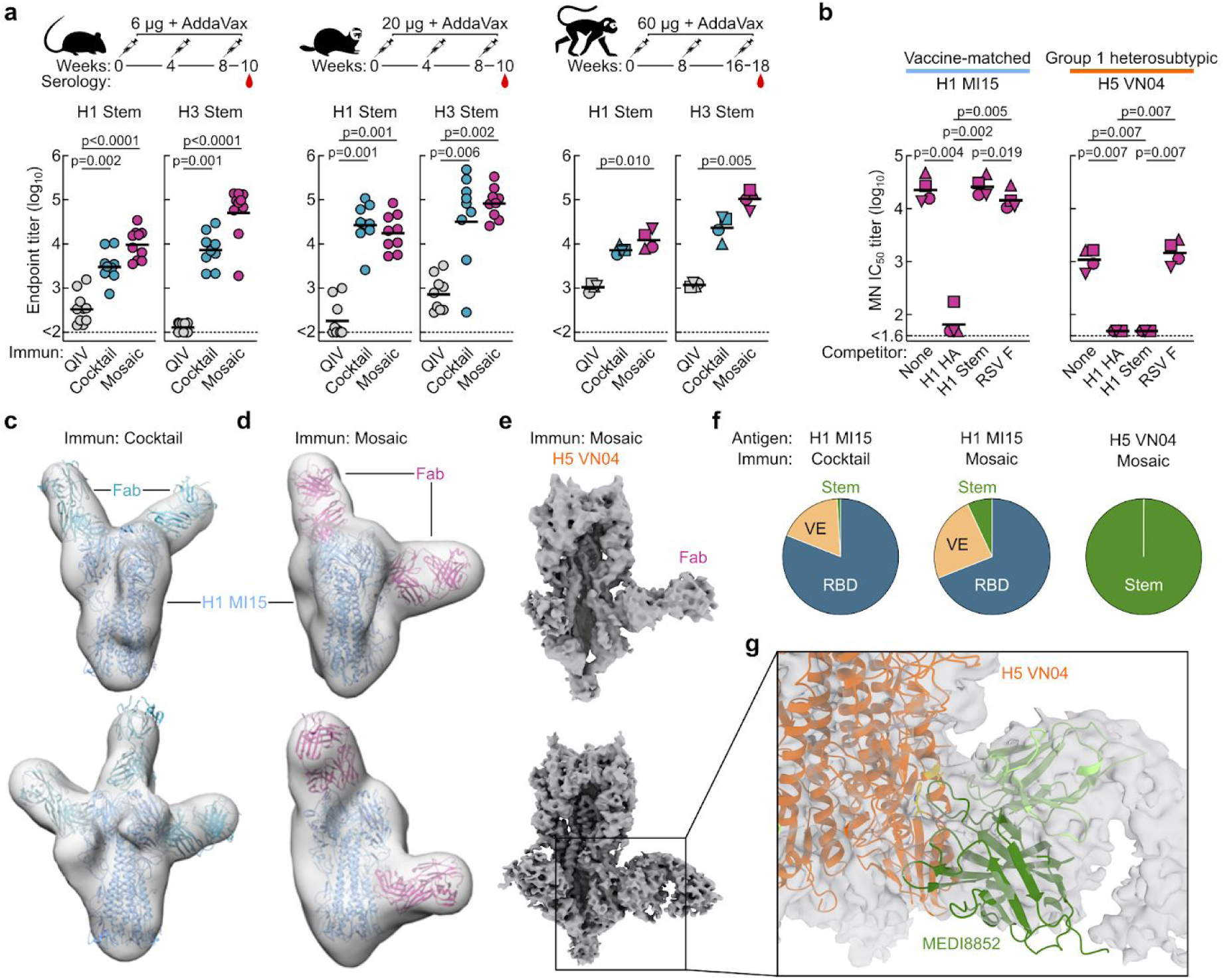
Molecular basis for nanoparticle immunogen-induced protection against heterosubtypic influenza viruses. **a**, Serum antibody titers to group 1 and 2 HA stem antigens in mice, ferrets, and NHPs. Immunization schemes are shown at the top of each panel. Antibody titers are expressed as endpoint dilutions. Each symbol represents an individual animal and the horizontal bar indicates the geometric mean of each group of BALB/cJ mice (*N* = 10), Finch ferrets (*N* = 9), and rhesus macaques (*N* = 4). Individual NHPs are identified by unique symbols. All animal immunization experiments except for NHP were performed at least twice and representative data are shown. **b**, Serum microneutralization activity in the presence of competitor proteins. Immune sera were evaluated directly or pre-incubated with either MI15 (H1N1) HA, CA09 (H1N1) stem HA, or irrelevant RSV F proteins prior to evaluation of MI15 (H1N1) and VN04 (H5N1) neutralization. Microneutralization titers are reported as half maximal inhibitory dilution (IC_50_). Statistical analysis was performed using nonparametric Kruskal–Wallis test with Dunn’s multiple comparisons. All animal experiments except for NHP were performed at least twice and representative data are shown. **c-d**, Selected EM reconstructions of negatively stained vaccine-matched H1 HA-foldon in complex with polyclonal antibody Fab fragments elicited by **c**, qsCocktail-I53_dn5 and **d**, qsMosaic-I53_dn5. The coordinates of an H1 HA crystal structure (PDB 1RUZ) and a Fab fragment (PDB 3GBN) were fitted into the EM densities. Light blue ribbons, H1 HA; cyan ribbons, polyclonal Fabs elicited by qsCocktail-I53_dn5; magenta ribbons, polyclonal Fabs elicited by qsMosaic-I53_dn5. **e**, Two independent cryo-EM reconstructions of heterosubtypic H5 HA-foldon in complex with polyclonal antibody Fab fragments elicited by qsMosaic-I53_dn5. Note the nearly orthogonal angles of approach adopted by the Fabs in the two reconstructions. **f**, Frequency of HA/Fab complexes observed by cryo-EM containing Fab fragments bound to RBD, VE, or stem domains. Single complexes containing Fabs of multiple specificities were counted once against each specificity. **g**, Close-up view showing that one of the dominant antibody recognition modes resembles that of MEDI8852 by forming putative contacts with helix A and the fusion peptide. The coordinates of an H5 HA crystal structure in complex with MEDI8852 Fab (PDB 5JW4) were fitted into the EM densities, with side-chains of Trp21_HA2_ and Ile45_HA2_ shown in gold.

We directly visualized the epitopes recognized by nanoparticle immunogen-elicited antibodies in individual NHPs using recently developed methods for single-particle EM analysis of polyclonal antibody responses^54,55^. We found that the polyclonal response to vaccine-matched H1 HA comprises at least three classes of antibodies targeting the receptor binding domain (RBD), the vestigial esterase (VE) domain, and the stem (Fig. 5c,d and Extended Data Figs. 8 and 9). While each class included multiple specificities recognizing distinct epitopes or having different angles of approach, the vast majority of complexes we visualized contained antibodies recognizing the head domain of H1 HA (Fig. 5f). We also carried out single-particle cryo-EM analysis of H5 HA in complex with polyclonal Fab fragments prepared from antibodies elicited by qsMosaic-I53_dn5. Strikingly, we saw only stem-directed antibodies bound to the H5 HA trimer, unambiguously demonstrating recognition of this conserved supersite (Fig. 5 e-g). 3D classification of the data revealed H5 HA trimers with polyclonal stem-directed Fabs bound to one, two, or all three HA subunits, from which we obtained 3D reconstructions with average resolutions ranging from 3.6 to 4.1 Å (Fig. 5e and Extended Data Fig. 10). Although the resolution of the Fab components is much lower than the nominal resolution due to polyclonal heterogeneity, the maps clearly indicate the existence of multiple vaccine-elicited antibody classes featuring distinct angles of approach to the epitope defined by the HA fusion peptide, helix A, and the hydrophobic groove surrounding Trp21_HA2_. The data suggest at least one common class of vaccine-elicited antibodies recognizes this epitope in a manner reminiscent of the human bnAbs MEDI8852 and 56.a.09, which both belong to the V_H_6-1+D_H_3-3 class of multi-donor human bnAbs^56,57^ (Fig. 5g). These findings corroborate our serological analyses and establish that although most vaccine-elicited antibodies recognize the vaccine-matched HA head, the heterosubtypic protection we observe likely derives from the simultaneous elicitation of stem-targeted bnAbs.

## Discussion

The manifest consequences of the SARS-CoV-2 pandemic emphasize the importance of efforts to mitigate the next influenza pandemic by establishing universal influenza immunity. We have developed nanoparticle vaccines that elicit potent vaccine-matched HAI activity as well as robust stem-directed antibodies against distantly related—including heterosubtypic—viruses in multiple animal models. While multiple next-generation influenza vaccine concepts have been reported to enhance either stem-directed responses^26–29,32,58^ or the potency and breadth of HAI within specific subtypes^34,35,59,60^, simultaneously inducing protective levels of both types of responses has not previously been achieved. Since both HAI and stem-directed antibodies have been shown as independent immune correlates of protection against influenza infection in humans^61^, a vaccine candidate capable of eliciting both would have advantages over approaches eliciting one or the other. The broad, antibody-mediated protection conferred by qsCocktail-I53_dn5 and qsMosaic-I53_dn5 suggests that they may be able to provide consistent year-to-year protection against seasonal influenza viruses, even in the event of antigenic mismatches in the hypervariable head domain. In contrast to universal vaccine approaches targeting only the HA stem, eliciting HAI responses against seasonal influenza strains with nanoparticle vaccines displaying full-length HAs justifies head-to-head evaluation against conventional licensed vaccines, which will facilitate clinical development. These unique features make our nanoparticle immunogens attractive candidates for clinical evaluation as supraseasonal vaccines^12^ that may eventually replace current seasonal vaccines.

Defining the immunological or structural basis for the singular breadth elicited by our nanoparticle vaccines will require further investigation. The induction of protective levels of stem-directed antibodies by both qsCocktail-I53_dn5 and qsMosaic-I53_dn5 despite the presence of the immunodominant head domain—but not by QIV or a non-assembling control immunogen—suggests that some aspect of HA presentation on the I53_dn5 scaffold alters the intrinsic immunodominance hierarchy of HA^23,62^. Several potential factors may contribute to this effect, such as the geometric relationship between neighboring HAs on the nanoparticle surface^33,42,45^, preferential antigen uptake or retention kinetics by macrophages and follicular dendritic cells^63^, or the presence of T helper epitopes within the subunits of the I53_dn5 nanoparticle^64–66^. Intriguingly, across our many *in vivo* studies, qsMosaic-I53_dn5 subtly, yet consistently, outperformed qsCocktail-I53_dn5 by most measures, including protective efficacy. Additional studies will be required to determine if this observation holds for mosaic and cocktail immunogens displaying different combinations of HAs and antigens other than HA.

We demonstrated the apparent epitope and angle of approach of a common class of vaccine-elicited, cross-reactive, polyclonal NHP antibodies are remarkably similar to human multi-donor V_H_6-1+D_H_3-3–class antibodies^56,57^, the broadest and most potent stem-directed human bnAbs^51^. Precursors of this class of bnAb are in theory found in ∼99% of the human population^67^ and can be amplified by heterologous stimuli such as pandemic influenza vaccines^57,68^. While this observation is encouraging, it remains to be seen whether more frequently observed stem-directed antibodies, such as those derived from V_H_1-69 genes, interfere with the elicitation of the broader and more potent V_H_6-1+D_H_3-3 antibodies in humans upon immunization with our nanoparticle vaccines, or whether other forms of pre-existing influenza immunity would alter the breadth of responses.

Our work establishes *in vitro* assembly as a key advantage of designed two-component nanoparticles^38,39,42^, as it enables facile and scalable co-display of multiple antigenic variants in stoichiometrically-controlled ratios. This property could be used to generate nanoparticle vaccine candidates co-displaying additional combinations of HAs for specific purposes. For example, incorporating additional HAs from co-circulating H3N2 or influenza B viruses could consolidate protective immunity against drifted seasonal strains. Including HAs from non-circulating subtypes, such as H5N1 and H7N9, may yield pandemic preparedness vaccines that provide broad protection against known and unknown pandemic threats^12^. While identifying the combinations and ratios of displayed HAs that elicit the broadest and most potent protective immunity will require further investigation, our data suggest that even displaying the four HAs of current seasonal influenza vaccines on I53_dn5, as we have done here, may provide moderate protection to unforeseen pandemic influenza viruses, which could provide the time needed for vaccine manufacturing in the event of an emerging threat^12^. The modularity of designed two-component nanoparticles could be a powerful tool in the development of next-generation influenza vaccines.

Motivated by the data presented here, a variant of qsMosaic-I53_dn5 with updated HA antigens is being produced under current Good Manufacturing Practice (cGMP) conditions for a planned Phase I clinical trial. Data from this trial should address several key questions raised by the current study, such as whether the protective stem-directed responses we observed in naïve animals can also be elicited in humans in the face of preexisting immunity. While animal models can be devised to investigate some aspects of this question (reviewed in ref. 69), human clinical trials will be needed to assess the impact of complex and individualized influenza exposure histories and immunological imprinting^70,71^. Detailed serological analyses in human subjects will also define the potential for qsMosaic-I53_dn5 to provide multi-season protection.

Understanding the mechanistic basis responsible for the exceptionally broad immune responses elicited by the nanoparticle immunogens described here may bring us a step closer to the ultimate goal of developing a universal influenza vaccine^72^. More generally, our work suggests that precisely designed two-component nanoparticles may be useful for generating vaccines that induce broadly protective immunity against other rapidly evolving pathogens and pandemic threats.

## Acknowledgments

The authors thank K. Foulds, A. Noe, S.-F. Kao, V. Ficca, N. Nji, D. Flebbe, and E. McCarthy (VRC) for help with nonhuman primate experiments; A. Taylor, H. Bao, C. Chiedi, M. Dillon, L. Gilman, and G. Sarbador, E. McCarthy, J.-P. Todd, Diana Scorpio (VRC) for help with mouse, ferret, and nonhuman primate experiments; H. Andersen, N. Jones, and G. Patel (Bioqual) for help with influenza challenge studies; R. Webby (St. Jude Children’s Research Hospital) for providing influenza reverse genetics plasmids; Y. Tsybovsky and T. Stephens (Frederick National Laboratory for Cancer Research) for initial EM screening; A. Reers and P. Myler (Seattle Children’s Research Institute) for assistance with protein production, and members of the King laboratory and the Influenza Program at the VRC for comments on the manuscript. This study was supported by the intramural research program of the Vaccine Research Center, National Institute of Allergy and Infectious Diseases, National Institutes of Health (M.K. and B.S.G.); a generous gift from the Open Philanthropy Project (D.B. and N.P.K.); a generous gift from the Audacious Project (D.B. and N.P.K.); the National Institute of General Medical Sciences (R01GM120553, D.V.); the National Institute of Allergy and Infectious Diseases (HHSN272201700059C to DV); a Pew Biomedical Scholars Award (D.V.); an Investigators in the Pathogenesis of Infectious Disease Award from the Burroughs Wellcome Fund (D.V.); and the University of Washington Arnold and Mabel Beckman cryo-EM center.

## Author Contributions

Conceptualization: B.S.G., N.P.K., M.K.; Modeling and design: D.E., G.U., N.P.K., M.K.; Formal Analysis: S.B.B., D.E., R.A.G., Y.J.P., O.A., M.J.W., S.K., K.K.L., M.G., D.V., N.P.K., M.K.; Investigation: S.B.B., D.E., R.A.G., G.B.H., A.C., Y.J.P., O.A., S.M.M., R.R., M.M., D.P., N.M., L.C., M.J.W., S.K., K.K.L., M.G., D.V., N.P.K., M.K.; Resources: A.C., G.U., L.S., D.B.; Writing – Original Draft: S.B.B., D.E., D.V., N.P.K., M.K.; Writing – Review & Editing: All authors; Visualization: S.B.B., D.E., Y.J.P., D.V., N.P.K., M.K.; Supervision: K.K.L., M.G., J.R.M., D.V., B.S.G., N.P.K., M.K.; Project Administration: M.C.C.; Funding Acquisition: L.S., D.V., J.R.M., B.S.G., D.B., N.P.K.

## Correspondence and requests for materials

should be addressed to B.S.G, N.P.K., and M.K. Influenza reverse genetics plasmids were provided by the St. Jude Children’s Research Hospital under a material transfer agreement with the NIH. Requests for these reagents should be made to the St. Jude Children’s Research Hospital.

## Methods

### Gene synthesis and vector construction

Plasmids for expression of the I53_dn5A pentamer were prepared in pET29b as previously described^42^. Genes for expression of HA fusions to nanoparticle trimeric components were codon optimized for expression in human cells and cloned into the CMV/R (VRC 8400) mammalian expression vector by Genscript. All HA fusions to the I53_dn5B trimer contained full-length HA ectodomains including native secretion signals, and the H1 and H3 HAs contained an additional mutation (Y98F) to knock out sialic acid binding to facilitate expression and purification^73^. HA ectodomain sequences preceded a short linker sequence followed by the I53_dn5B trimer sequence with a C-terminal flexible linker, WELQut protease recognition sequence, and a hexa-histidine tag. The amino acid sequences for all proteins used in this study are provided in Supplementary Table 1.

### Protein expression and purification

All HA-I53_dn5B trimers, as well as mAbs CR6261 (ref. 74), 5J8 (ref. 75), CR8020 (ref. 76), F005-126 (ref. 77), F045-092 (ref. 78), MEDI8552 (ref. 56), FI6v3 (ref. 53), CR9114 and CR8071 (ref. 79), CT149 (ref. 80), D25 (ref. 81), and MPE8 (ref. 82) were expressed in Expi293F cells (ThermoFisher Scientific) by transient transfection using PEI MAX (Polysciences) or ExpiFectamine™ 293 (ThermoFisher Scientific). mAbs were purified by protein A affinity chromatography using established methods. Recombinant HA ectodomain trimers fused to T4 fibritin foldon were produced and purified as described previously^34^. The protein-containing supernatants from cells expressing HA-I53_dn5B fusion proteins were further clarified by vacuum filtration (0.22 µm, Millipore Sigma). Prior to immobilized metal affinity chromatography, a background of 50 mM Tris pH 8.0 and 350 mM NaCl was added to the clarified supernatant using concentrated solutions of 1 M Tris pH 8.0 and 5 M NaCl, respectively. For each liter of supernatant, 4 mL of Ni^2+^ Sepharose Excel resin (GE) was rinsed into PBS using a gravity column and then added to the supernatant, followed by overnight shaking at 4°C. The resin was collected 16-24 h later using a gravity column, then washed twice with 50 mM Tris pH 8.0, 500 mM NaCl, 30 mM imidazole prior to elution of His-tagged protein using 50 mM Tris pH 8.0, 500 mM NaCl, 300 mM imidazole. Eluates were concentrated and applied to a HiLoad 16/600 Superdex 200 pg column or a Superdex 200 Increase 10/300 GL column pre-equilibrated with PBS for preparative size exclusion chromatography. Peaks corresponding to trimeric species were identified based on elution volume and SDS-PAGE (both reducing and non-reducing) of elution fractions. Fractions containing pure HA-I53_dn5B were pooled and the protein quantified using UV/vis spectroscopy. Purified protein was either stored at 4°C until use or flash-frozen in liquid nitrogen and stored at −80°C.

Single colonies *of E. coli* cells transformed with plasmid encoding the I53_dn5A pentamer were picked and grown in TB medium with 50 µg L^-1^ kanamycin at 37°C overnight. Subsequently, liquid cultures were diluted 1:40 in TB medium and grown at 37°C until OD_600_ reached 0.5-0.8. Isopropyl-thio-β-D-galactopyranoside (IPTG) was then added to a concentration of 1 mM and the growth temperature reduced to 18°C to induce protein expression, or cultures were left at 37°C for an additional 5 h before lowering the temperature to 18°C leading to auto-induction by virtue of galactose in the media according to the Studier protocols^83^. Expression proceeded for 20 h at 18°C, at which point the cell cultures were harvested by centrifugation. Cell pellets were resuspended in 50 mM Tris pH 8.0, 250 mM NaCl, 20 mM imidazole, 1 mM dithiothreitol (DTT), 0.1 mg ml^-1^ DNase, 0.1 mg ml^-1^ RNase, and EDTA-free protease inhibitors (Pierce) and lysed by sonication or microfluidization. I53_dn5A protein was purified from the soluble fraction of cell lysates by immobilized metal affinity chromatography using HisTrap HP columns (GE). After application of clarified cell lysate supernatants, the column was washed with 20 column volumes of 50 mM Tris pH 8.0, 250 mM NaCl, 20 mM imidazole, 1 mM DTT. I53_dn5A was eluted using a linear gradient of imidazole up to a final concentration of 500 mM. Protein was concentrated to 1 mL, and 3-[(3-Cholamidopropyl)dimethylammonio]-1-propanesulfonate (CHAPS) was added to 0.75% (w/v) to remove endotoxin. The protein was sterile-filtered at 0.22 μm and purified by preparative SEC using a Superdex 200 Increase 10/300 GL equilibrated in 25 mM Tris pH 8.0, 150 mM NaCl, 5% glycerol. The peak corresponding to the pentamer was identified based on elution volume and SDS-PAGE (both reducing and non-reducing) of eluted fractions. Fractions containing pure I53_dn5A pentamer were pooled and the protein quantified using UV/vis spectroscopy. The samples were confirmed to be low in endotoxin (<100 EU mg^-1^) using the Limulus Amebocyte Lysate (LAL) assay (Charles River), then flash-frozen in liquid nitrogen and stored at −80°C within 6 hours of purification to prevent oxidation of cysteines.

### *In vitro* assembly and purification of nanoparticle immunogens

To assemble nanoparticle immunogens bearing multiple copies of single HA antigens (e.g., H1-I53_dn5), individual HA-bearing trimeric components were mixed with pentameric I53_dn5A at a molar ratio of 1:1 (subunit:subunit) at concentrations ranging from 15-40 μM (subunit) by pipetting. Assembly reactions were prepared at room temperature and incubated for 30 min before purification by SEC on a Superose 6 Increase 10/300 GL. The nanoparticle immunogens eluted at the void volume of the column. Fractions were analyzed by SDS-PAGE (both reducing and non-reducing) before pooling and sterile filtering at 0.22 μm.

For H1- and H3-bearing components, the assembly reactions consisted of pentameric components and HA-bearing trimeric components buffered in either PBS or 25 mM Tris pH 8.0, 150 mM NaCl, 5% glycerol. After assembly and incubation, the samples were centrifuged for 10 min at 14,000 rpm at 4°C and the nanoparticle immunogens purified by SEC using a Superose 6 Increase 10/300 GL column pre-equilibrated with 25 mM Tris pH 8.0, 150 mM NaCl, 5% glycerol.

For B/Vic- and B/Yam-bearing components, half of the volume of the assembly solution consisted of an additional buffer solution with high ionic strength to maintain nanoparticle immunogen solubility. The solutions used were 25 mM Tris pH 8.0, 1.85 M NaCl, 5% glycerol for B/Vic and 25 mM Tris pH 8.0, 3.85 M NaCl, 5% glycerol for B/Yam, which respectively brought NaCl in the assembly reactions to approximately 1 M and 2 M. In these cases the HA-bearing trimeric component was first added to the high-salt buffer prior to addition of the pentameric component. After assembly and incubation, the samples were centrifuged for 10 min at 14,000 rpm at room temperature and the nanoparticle immunogens purified by SEC using a Superose 6 Increase 10/300 GL column pre-equilibrated with either 25 mM Tris pH 8.0, 1 M NaCl, 5% glycerol for B/Vic-I53_dn5 or 25 mM Tris pH 8.0, 2 M NaCl, 5% glycerol for B/Yam-I53_dn5.

For mosaic nanoparticles with equal amounts of each seasonal HA (qsMosaic-I53_dn5), all four HA-bearing trimeric components (in PBS) were first mixed in equimolar amounts. Tris pH 8.0, 1.85 M NaCl, 5% glycerol was added such that the final NaCl in the *in vitro* assembly reaction would be 1 M. The pentameric component was added and the solution was mixed vigorously by pipetting. After assembly and incubation, the samples were centrifuged for 10 min at 14,000 rpm at 4°C and the nanoparticle immunogens purified by SEC using a Superose 6 Increase 10/300 GL column pre-equilibrated with 25 mM Tris pH 8.0, 150 mM NaCl, 5% glycerol.

After purification and evaluation of nanoparticle immunogen quality by SDS-PAGE, UV/vis spectroscopy, negative stain EM, DLS, and LAL assay (<100 EU mg^-1^), samples were flash-frozen in liquid nitrogen and stored at −80°C.

### Dynamic light scattering

Light scattering analysis was conducted using an UNcle (UNchained Labs) at 25°C. For each sample, 10 acquisitions (5 s per acquisition) were obtained using auto-attenuation of the laser. Increased viscosity due to the inclusion of 5% glycerol in the H1-I53_dn5, H3-I53_dn5, B/Yam-I53_dn5, B/Vic-I53_dn5, qsMosaic-I53_dn5 and I53_dn5 nanoparticles was accounted for in the software.

### Negative stain and cryo-electron microscopy of immunogens

To image nanoparticles and non-assembling immunogens by negative stain EM, protein samples were diluted to 0.020-0.075 mg ml^-1^ in 25 mM Tris pH 8.0 with NaCl concentrations ranging from 0.15-2 M. 300 mesh copper grids (Ted Pella) were glow discharged immediately before use. Six μL of sample was applied to the grid for 1 min, then briefly dipped in a droplet of water before blotting away excess liquid with Whatman No. 1 filter paper. Grids were stained with 6 μL of 0.75% (w/v) uranyl formate stain, immediately blotting away excess, then stained again with another 6 μL for 30 s. Grids were imaged on a Morgagni transmission electron microscope with a Gatan camera, and Gatan Digital Micrograph software was used to take images.

To obtain a cryo-EM single particle reconstruction of the H1-I53_dn5 nanoparticle, 3 µL of 0.7 mg ml^-1^ H1-I53_dn5 was loaded onto a freshly glow-discharged (30 s at 20 mA) Protochips C-flat grid (2.0 μm hole, 200 mesh) by multiple blotting strategy prior to plunge freezing using a vitrobot Mark IV (ThermoFisher Scientific) using a blot force of 0 and 7 second blot time at 100% humidity and 25°C. Data was collected using the Leginon software on an FEI Titan Krios transmission electron microscope, equipped with a Gatan K2 Summit direct electron detector and Gatan Quantum GIF energy filter, operated in zero-loss mode with a slit width of 20 eV. The dose rate was adjusted to 8 counts pixel^-1^ s^-1^, and each movie was acquired in counting mode fractionated in 50 frames of 200 ms. 803 micrographs were collected in a single session with a defocus range between −1.2 μm and −2.5 μm. Movie frame alignment and estimation of the microscope contrast-transfer function parameters were carried out using Warp^84^. A total of 2,000 particles were manually picked and 2D classifications were performed in RELION 3.0 (ref. 85). Five representative class averaged images were selected as references for automatic particle picking. 2D and 3D classification and 3D refinements were performed using RELION 3.0. A total of 20,827 particles of the best class were selected and subjected to 3D refinement. We subsequently implemented a previously reported localized reconstruction strategy^47^ to determine a reconstruction of the H1 MI15 HA at 3.3 Å resolution using Relion. Reported resolutions are based on the gold-standard Fourier shell correlation (FSC) of 0.143 criterion and Fourier shell correlation curves were corrected for the effects of soft masking by high-resolution noise substitution^86^.

### Immunoprecipitation

qsCocktail-I53_dn5 and qsMosaic-I53_dn5 samples were mixed with either MPE8 (anti-RSV F) or 5J8 (anti-H1) to final concentrations of 4 μM (each subunit) of immunogen and 0.20 mg ml^-1^ of mAb. The final buffers contained 2 M NaCl and 0.05% Tween-20 in addition to other buffering agents. The solution was allowed to mix at room temperature for 1 h and then added to recently washed and dried magnetic protein G Dynabeads (Thermo Fisher). The mixture was incubated for 1.5 h and resin was separated from the supernatant magnetically. Beads were washed twice and returned to the same volume used during the binding process, then heated to 95°C for 10 min in the presence of SDS loading dye to detach bound protein. Proteins were then analyzed by SDS-PAGE.

### Antigenic characterization

ELISA was used to measure binding of H1-I53_dn5, H3-I53_dn5, B/Yam-I53_dn5, B/Vic-I53_dn5, and qsMosaic-I53_dn5 nanoparticles to mAbs CR6261, 5J8, CR8020, F005-126, MEDI8852, FI6v3, CR9114, CR8071, and D25 (anti-RSV F). 96-well plates were coated with 2 µmol ml-1 nanoparticles (0.1 ml well^-1^) and incubated at 4°C overnight. Plates were then blocked with PBS containing 5% skim milk at 37°C for 30 min. mAbs were serially diluted in four-fold steps and added to the wells for 1 h. Horseradish peroxidase (HRP)-conjugated anti-human or anti-mouse IgG (Southern Biotech) was added and incubated at 37°C for 30 min. The wells were developed with 3,3′,5′,5-tetramethylbenzidine (TMB) substrate (KPL), and the reactions were stopped by adding 1 M H_2_SO_4_ before measuring absorbance at 450 nm with a Spectramax Paradigm plate reader (Molecular Devices).

### Mass spectrometry quantification of HA content in qsMosaic-I53_dn5

Label-free quantitation was performed by peptide mass spectrometry to determine the relative abundance of each HA present in the mosaic nanoparticle samples. Each mosaic nanoparticle, either before or after SEC purification, along with a standard mixture of each purified HA-I53_dn5B fusion protein at equimolar concentrations (1:1:1:1), was denatured and reduced using guanidine hydrochloride and DTT. Samples were then alkylated with iodoacetamide, deglycosylated with N-glycanase (New England Biolabs), and digested overnight with LysC protease (ThermoFisher scientific). LC-MS was performed using a Waters Acquity UPLC coupled to a Thermo LTQ-OT using data-dependent acquisition. Peptides were resolved over a Waters CSH C18 1.7 μm, 2.1×100 mm column with a linear gradient from 3% to 40% B over 30 minutes (A: 0.1% formic acid; B: acetonitrile with 0.1% formic acid). Peptides were identified from MS/MS data using Protein Prospector using a score cutoff of 15 (http://prospector.ucsf.edu/). Due to the high sequence identity between the HA constructs, only four peptides unique to each specific HA were observed that could be used for label-free quantitation. The integrated peak areas for these peptides relative to the areas from an equimolar mixture of each HA were used to estimate the total abundance of each HA within the mosaic nanoparticle samples (Supplementary Table 3).

### Hydrogen-deuterium exchange mass spectrometry

For each timepoint, 30 pmol of H1 HA-foldon and H1-I53_dn5 were incubated in deuterated buffer (85% D_2_O, pH* 7.4) for 3; 60; 1,800; or 72,000 s at room temperature and subsequently mixed with an equal volume of ice-cold quench buffer (4 M urea, 200 mM tris(2-chlorethyl) phosphate (TCEP), 0.2% formic acid) to a final pH* of 2.5. Samples were immediately frozen in liquid nitrogen and stored at 80°C until analysis. Fully deuterated samples were prepared by digesting 30 pmol of undeuterated H1-foldon over a pepsin column, followed by concentration under vacuum, resuspension in deuterated buffer at 65°C for 1 h, then quenching and freezing. Zero timepoint samples were prepared as described previously^87^. Inline pepsin digestion was performed and analyzed by LC-IMS-MS utilizing a Waters Synapt G2-Si Q-TOF mass spectrometer as previously described^87^. Deuterium uptake analysis was performed using HD-Examiner (Sierra Analytics) followed by HX-Express v3.13 (refs. 88,89). The percent exchange was normalized to the zero timepoint and fully deuterated reference samples. Internal exchange standards (Pro-Pro-Pro-Ile [PPPI] and Pro-Pro-Pro-Phe [PPPF]) were included in each reaction to ensure that conditions were consistent throughout all of the labeling reactions.

### Animal experiments

All animal experiments were reviewed and approved by the Institutional Animal Care and Use Committee of the VRC, NIAID, NIH. All animals were housed and cared for in accordance with local, state, federal, and institutional policies of NIH and American Association for Accreditation of Laboratory Animal Care.

### Immunization and challenge studies

Throughout our studies, we matched the total protein dose of each nanoparticle immunogen to the HA content of QIV. Because the HA antigens make up approximately 62% of the total peptidic mass of the nanoparticle immunogens, animals immunized with commercial QIV received ∼1.6× as much HA as animals immunized with the nanoparticle immunogens. BALB/cJ mice (Jackson Laboratory) were immunized intramuscularly (i.m.) with 6 μg of commercial QIV 2017-2018, qsCocktail-I53_dn5, or qsMosaic-I53_dn5 in the presence or absence of AddaVax™ (InvivoGen) at weeks 0, 4, and 8. Formulated vaccines were given 50 μl into each hind leg. Serum samples were collected before and after each immunization and used for immunological assays. For the challenge studies, mice were infected intranasally with 10-20 times the 50% lethal dose (LD_5s0_) of H1N1, H5N1, H3N2, or H7N9 viruses (Supplementary Table 2) at Bioqual. The animals were monitored twice daily for development of clinical signs and weighed daily for 14 days. Any animals that had lost 20% or more of their initial body weight were euthanized. Finch ferrets (*Mustela putorius*) were immunized i.m. with 20 μg of commercial QIV 2017-2018, qsCocktail-I53_dn5, or qsMosaic-I53_dn5 with AddaVax three times at weeks 0, 4, and 8. Immunogens were formulated in 500 μL per ferret and injected into limbs. Serum samples were collected periodically before and after immunization and used for immunological assays. For the challenge studies, ferrets were infected intranasally with 10-20 times the 50% lethal dose (LD_50_) of H5N1 or H7N9 viruses (Supplementary Table 2) at Bioqual. Clinical signs of infection, weight, and body temperatures were recorded twice daily for 14 days. Activity scores were assigned as follows: 0, alert and playful; 1, alert but playful only when stimulated; 2, alert, but not playful when stimulated; and 3, neither alert nor playful when stimulated. Ferrets that showed signs of severe disease (prolonged fever, diarrhea; nasal discharge interfering with eating, drinking or breathing; severe lethargy; or neurological signs) or that had >20% weight loss were euthanized immediately. Rhesus macaques (*Macaca mulatta*) were immunized i.m. with 60 μg of commercial QIV 2017-2018, qsCocktail-I53_dn5, or qsMosaic-I53_dn5 with AddaVax three times at weeks 0, 8, and 16. Immunogens were prepared in 1.0 mL volumes per NHP and injected into limbs. Serum samples were collected periodically before and after immunization and used for immunological assays.

### ELISA

ELISA was used to measure HA-specific IgG levels in immune sera. The plates were coated with 2 μg ml^-1^ of recombinant HA-foldon proteins (Supplementary Table 1) and incubated at 4°C overnight. Plates were then blocked with PBS containing 5% skim milk at 37°C for 1 h. mAbs and immune sera were serially diluted in four-fold steps and added to the wells for 1 h. Horseradish peroxidase (HRP)-conjugated anti-human or anti-mouse IgG (Southern Biotech) was added and incubated at 37°C for 1 h. The wells were developed with 3,3′,5′,5-tetramethylbenzidine (TMB) substrate (KPL), and the reactions were stopped by adding 1 M H_2_SO_4_ before measuring absorbance at 450 nm with a Spectramax Paradigm plate reader (Molecular Devices).

### Reporter-based microneutralization assay

All reporter viruses were prepared as described previously^51^. Briefly, all H1N1 and H3N2 viruses were made with a modified PB1 segment expressing the TdKatushka reporter gene (R3ΔPB1) and propagated in MDCK-SIAT-PB1 cells, while H5N1 reporter virus was made with a modified HA segment expressing the reporter (R3ΔHA) and produced in cells stably expressing H5 HA. Replication-restricted reporter influenza viruses encoding influenza B HA and NA coding regions were rescued using plasmids expressing the open reading frames of influenza B HA and NA genes flanked by genome packaging signals of influenza A HA^90^ and NA segments^91^, respectively. These viruses have a PB1 segment modified to express the TdKatushka2 reporter gene and encode the internal genes of influenza A (A/WSN/1933, H1N1) virus. Rescued viruses were propagated in MDCK-SIAT1-PB1 in the presence of TPCK-trypsin (1 μg mL^-1^, Sigma) at 34°C. Virus stocks were stored at −80°C. Mouse sera were treated with receptor destroying enzyme (RDE II; Denka Seiken) and heat-inactivated before use in neutralization assays. Immune sera or mAbs were serially diluted and incubated for 1 h at 37°C with pre-titrated viruses (Supplementary Table 2). Serum-virus mixtures were then transferred to 96-well plates (PerkinElmer), and 1.0×10^4^ MDCK-SIAT1-PB1 cells^51,92^ were added into each well. After overnight incubation at 37°C, the number of fluorescent cells in each well was counted automatically using a Celigo image cytometer (Nexcelom Biosciences).

### Hemagglutination inhibition titer

HAI titer to vaccine-matched viruses were tested with immune sera. The reporter influenza viruses H1N1 MI15, B/Vic CO17, and B/Yam PH13 (Supplementary Table 2) were propagated in Madin-Darby canine kidney (MDCK) cells. Immune sera were treated with receptor-destroying enzyme (RDE II; Denka Seiken) before use in HAI assays. Immune sera and mAbs were serially diluted and incubated with reporter viruses and then incubated with 0.5% turkey red blood cells (Lampire Biological Laboratories) for 30 min at room temperature. Following the incubation period, the plates were analyzed to distinguish between agglutinated wells with diffuse reddish appearance and non-agglutinated wells containing a dark red pellet. The HAI titer of the sample was determined based on the well with the last agglutinated appearance, immediately before a pellet was observed.

### Passive transfer studies

To generate hyper-immune Ig for passive transfer, the immune serum samples from each NHP were diluted 1:50 with PBS, added to protein A columns, and incubated overnight at 4°C. After washing the columns briefly, captured antibodies were eluted with low-pH IgG elution buffer (ThermoFisher Scientific) and the eluates were immediately neutralized by adding 1 M Tris-HCl (pH 8.0) to a final concentration of 100 mM. Purified polyclonal antibodies were dialyzed two times against PBS, concentrated to ∼20 mg ml-1 and stored at −80°C until use. BALB/cAnNHsd mice (Envigo) were given intraperitoneally 0.2 mg of FI6v3 (approximately 10 mg/kg) or 10 mg of purified polyclonal Ig from individual NHPs. 24 h later, the mice were infected intranasally with 10-fold the LD_50_ of H5N1 or H7N9 viruses (Supplementary Table 2) at Bioqual. The animals were monitored twice daily for development of clinical signs and weighed daily for 14 days. Any animals that lost 20% or more of their initial body weight were euthanized.

### Preparation of polyclonal immunoglobulin antigen-binding fragments

To generate polyclonal Fab fragments for epitope mapping, the immune serum samples from each NHP were diluted with PBS, applied to protein A columns, and incubated overnight at 4°C. After washing the columns briefly, captured antibodies were eluted with 0.1 M glycine pH 3.5, and the eluates were immediately neutralized by adding Tris-HCl (pH 8.0) to a final concentration of 50 mM. Purified IgG was buffer-exchanged into PBS and concentrated to approximately 25 mg ml^-1^, and 250 μL of 2× digestion buffer (40 mM sodium phosphate pH 6.5, 20 mM EDTA, 40 mM cysteine) was added. 500 μL of resuspended immobilized papain resin (ThermoFisher Scientific) freshly washed in 1× digestion buffer (20 mM sodium phosphate, 10 mM EDTA, 20 mM cysteine, pH 6.5) was further added, and samples were shaken for 5 h at 37°C. The supernatant was separated from resin and mixed with 1 mL of 20 mM Tris pH 8.0. Resin was washed twice with 500 μL of 20 mM Tris pH 8.0 and supernatants from the washes were pooled with the original supernatant to increase sample yield. Pooled supernatants were sterile-filtered at 0.22 μm and applied to protein A columns. Unbound fractions were pooled, concentrated to approximately 10 mg ml^-1^, and dialyzed twice against 25 mM Tris pH 8.0 to remove excess phosphates and cysteine prior to sample preparation for EM. Final samples were confirmed by SDS-PAGE, flash-frozen, and stored at −80°C.

### Electron microscopy polyclonal epitope mapping (EMPEM)

To prepare H1 HA/polyclonal Fab fragment complexes, 150-fold molar excesses of qsCocktail- or qsMosaic-I53_dn5-elicited antibody Fab fragments were incubated with H1 HA-foldon for 1 h at room temperature, and the complexes were purified on a Superdex 200 Increase 10/300 GL column. The purified complexes were adsorbed onto glow-discharged carbon-coated copper mesh grids for 60 s, stained with 2% uranyl formate for 30 s, and allowed to air dry. Grids were imaged using an FEI Tecnai Spirit 120 kV electron microscope equipped with a Gatan Ultrascan 4000 CCD Camera. The pixel size at the specimen level was 1.60 Å. Data collection was performed using Leginon^93^ with the majority of the data processing carried out in Appion^94^. 4,112 and 3,237 micrographs were collected for qsCocktail-I53_dn5- and qsMosaic-I53_dn5-elicited Fab/HA complexes respectively. The parameters of the contrast transfer function (CTF) were estimated using CTFFIND4 (ref. 95). All particles were picked in a reference-free manner using DoG Picker^96^. Reference-free 2D classification was used to select homogeneous subsets of particles using CryoSPARC^97^. 847,873 and 997,557 particles were subjected for 2D classification of qsCocktail-I53_dn5- and qsMosaic-I53_dn5-elicited Fab/HA complexes, respectively. During the 2D classification, 2D classes were visually inspected and particles from classes not showing clear structural features of Fab/HA complexes were discarded. The remaining particles were subsequently subjected to three rounds of ab initio 3D reconstructions and 3D classifications without any symmetry imposed using CryoSPARC. Only receptor binding domain, vestigial esterase domain, and stem-directed Abs were included in the calculations. Particles from these classes were separately subjected to 3D refinement using CryoSPARC. The head-binding Fabs of the different classes were similar, but most classes showed obvious asymmetric features. All 3D reconstructions were compared to three classes of structurally characterized anti-HA antibodies: (i) receptor binding domain-targeted Abs CH65 (PDB: 5UGY), C05 (PDB: 4FP8), F045-092 (PDB: 4O58), HC63 (PDB: 1KEN), 2G1 (PDB: 4HG4), 8M2 (PDB: 4HFU), 5J8 (PDB: 4M5Z), 1F1 (PDB: 4GXU), and S139/1 (PDB: 4GMS); (ii) vestigial esterase domain-targeted Abs H5M9 (PDB: 4MHJ) and CR8071 (PDB: 4FQJ); and (iii) stem-binding Abs C179 (PDB: 4HLZ), CR6261 (PDB: 3GBN), CR8043 (PDB: 4NM8), CR8020 (PDB: 3SDY), CR9114 (PDB: 4FQI), FI6v3 (PDB: 3ZTJ), MEDI8852 (PDB: 5JW4), and 39.29 (PDB: 4KVN). Single particles belonging to each 3D reconstruction were re-grouped into three and counted. Estimates of the fraction of particles containing receptor binding domain-, vestigial esterase domain-, and stem-binding Fabs were based on the number of particles clustered in each group. Particles containing Fabs bound to multiple sites were counted against each site.

H5 HA/polyclonal Fab fragment complexes were prepared and verified by negative stain electron microscopy as described above and then pooled and concentrated. 3 µL of 0.1 mg ml^-1^ H5 HA in complex with qsMosaic-I53_dn5-elicited antibody Fab fragments was loaded onto a freshly glow-discharged (30 s at 20 mA) 1.2/1.3 UltraFoil grid (300 mesh) with a thin layer of evaporated continuous carbon prior to plunge freezing using a vitrobot Mark IV (ThermoFisher Scientific) using a blot force of −1 and 2.5 second blot time at 100% humidity and 25°C. Data were acquired using the an FEI Titan Krios transmission electron microscope operated at 300 kV and equipped with a Gatan K2 Summit direct detector and Gatan Quantum GIF energy filter, operated in zero-loss mode with a slit width of 20 eV. Automated data collection was carried out using Leginon at a nominal magnification of 130,000× with a pixel size of 0.525 Å. The dose rate was adjusted to 8 counts/pixel/s, and each movie was acquired in super-resolution mode fractionated in 50 frames of 200 ms. 2,374 micrographs were collected using beam-image shift with a defocus range between −1.0 and −2.5 μm. Movie frame alignment, estimation of the microscope contrast-transfer function parameters, particle picking, and extraction were carried out using Warp. Particle images were extracted with a box size of 800 binned to 400 yielding a pixel size of 1.05 Å. Two rounds of reference-free 2D classification were performed using CryoSPARC to select well-defined particle images. These selected particles were subjected to 3D refinement in CryoSPARC. For beam tilt correction, the micrographs were grouped into beam tilt groups using beam-image shift values from Leginon. Beam tilt refinement was performed in Relion3.0. CTF refinement was used to refine per-particle defocus values. Particle images were subjected to the Bayesian polishing procedure implemented in Relion3.0. After determining a refined 3D structure, the particles were then subjected to 3D classification without refining angles and shifts using a soft mask on three Fab regions and with a tau value of 20. 3D refinements were carried out using non-uniform refinement along with per-particle defocus refinement in CryoSPARC. Local resolution estimation, filtering, and sharpening was carried out using CryoSPARC. Reported resolutions are based on the gold-standard Fourier shell correlation (FSC) of 0.143 criterion and Fourier shell correlation curves were corrected for the effects of soft masking by high-resolution noise substitution.

### Statistics and Reproducibility

Multi-group comparisons were performed using nonparametric Kruskal-Wallis test with Dunn’s post-hoc analysis in Prism 8 (GraphPad) unless mentioned otherwise. Differences were considered significant when P values were less than 0.05. Statistical methods and P value ranges can be found in the Figures and Figure legends.

## Data availability

All images and data were generated and analyzed by the authors, and will be made available by the corresponding authors (B.S.G., N.P.K., and M.K.) upon reasonable request. Uncropped images of all gels/blots are provided in Supplementary Figure 1. Maps for the single-particle reconstruction of the H1-I53_dn5 nanoparticle and the local reconstruction of the displayed H1 HA will be deposited in EMDB prior to publication.

