## Extended Data for "Elicitation of broadly protective immunity to influenza by multivalent hemagglutinin nanoparticle vaccines"

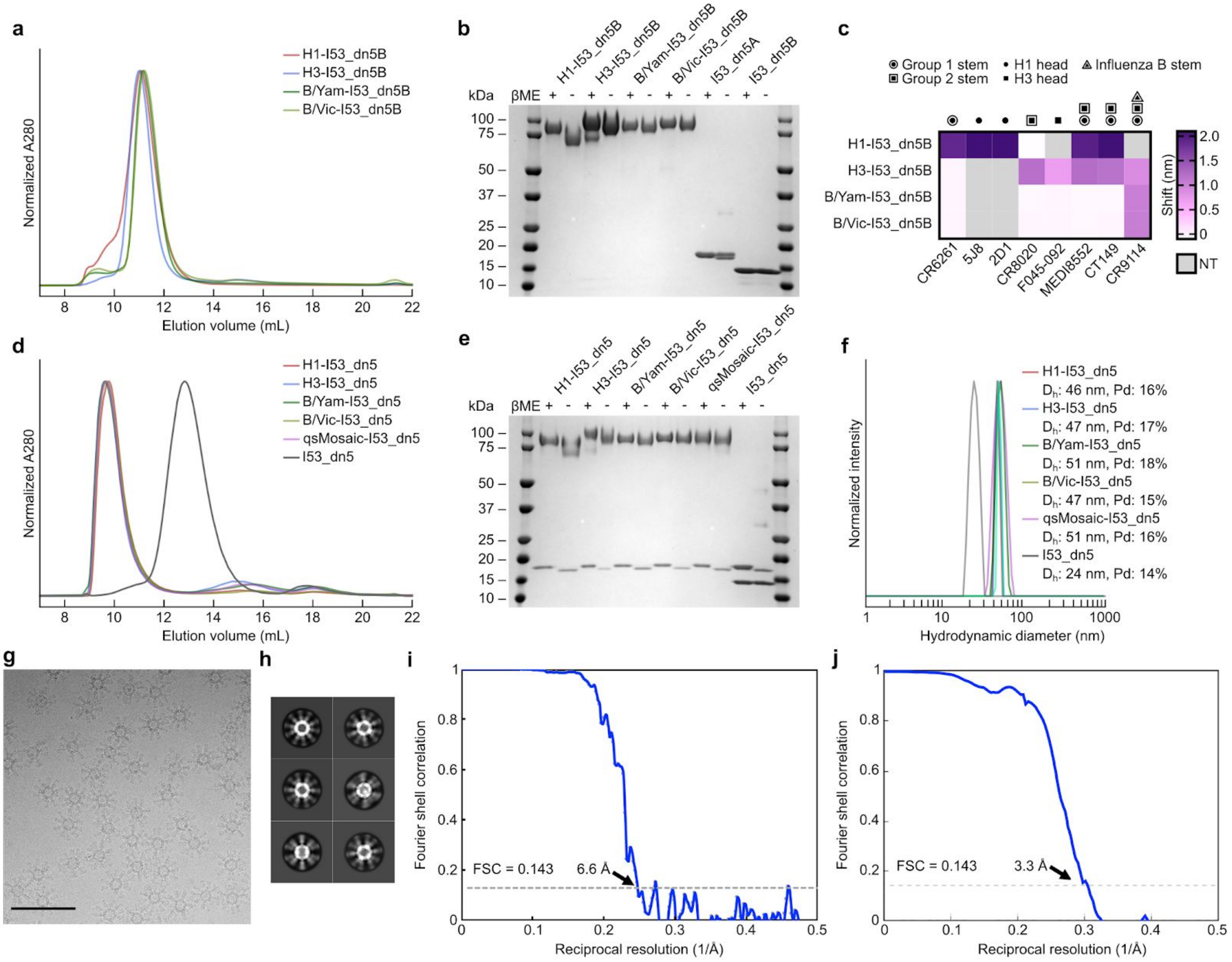


**Extended Data Fig. 1 | Production and characterization of HA-I53_dn5 components and nanoparticle immunogens. a,** SEC purification of seasonal HAs fused to I53_dn5B trimeric components, using a Superdex 200 Increase 10/300 GL column. **b,** Reducing and non-reducing SDS-PAGE of SEC-purified trimeric HA-I53_dn5B fusions, pentameric I53_dn5A component and I53_dn5B trimer lacking fused HA. **c,** Antigenic characterization of HA-I53_dn5B trimeric components by BLI. Symbols indicate the specificity of each mAb. NT, not tested. **d,** SEC purification of nanoparticle immunogens after *in vitro* assembly, including I53_dn5 lacking displayed antigen, using a Superose 6 Increase 10/300 GL column. The nanoparticle immunogens elute at the void volume of the column, while I53_dn5 is resolved. Residual, unassembled trimeric and pentameric components elute around 15 mL and 18 mL, respectively. **e,** Reducing and non-reducing SDS-PAGE of SEC-purified nanoparticle immunogens and I53_dn5. **f,** Dynamic light scattering (DLS) of SEC-purified nanoparticle immunogens, including I53_dn5. **g,** Representative electron micrograph of H1-I53_dn5 embedded in vitreous ice. Scale bar, 100 nm. **h,** 2D class averages obtained using single-particle cryo-EM. **i,** Gold-standard Fourier shell correlation curve for the H1-I53_dn5 density map presented in Fig. 1e. **j,** Gold-standard Fourier shell correlation curve for the local reconstruction of H1 MI15 presented in Fig. 1e.

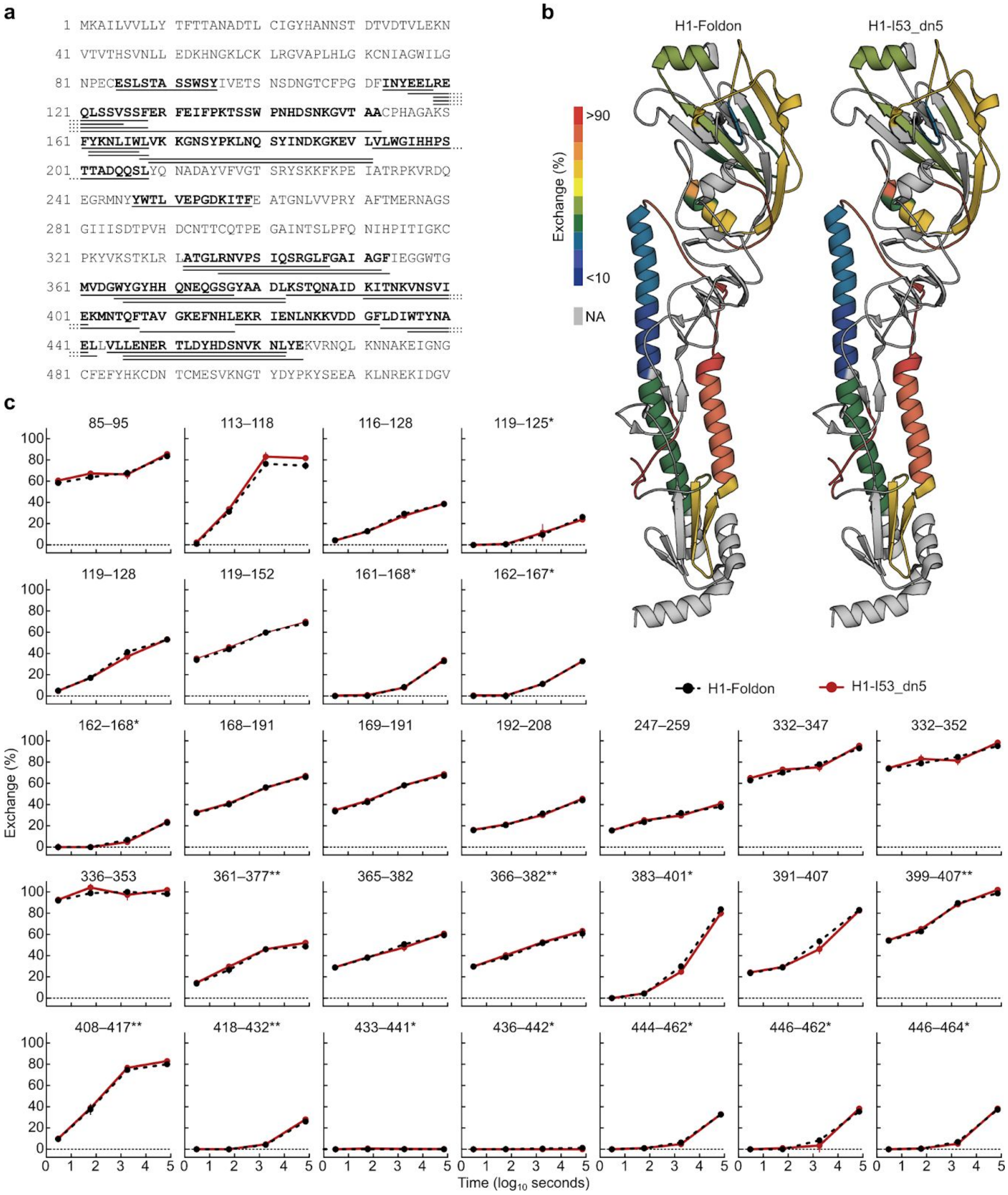


**Extended Data Fig. 2 | Hydrogen-deuterium exchange mass spectrometry (HDX-MS) of H1-foldon trimer and H1-I53_dn5 nanoparticle. a,** Amino acid sequence of H1 ectodomain expressed as a genetic fusion to both foldon and I53_dn5B. Underlined sequences correspond to peptides analyzed by HDX-MS. **b,** Hydrogen-deuterium exchange percentages after 20 h for both samples mapped onto the structure of H1 HA (PDB 3LZG). **c,** Kinetics of hydrogen-deuterium exchange for both samples at multiple timepoints up to 20 h. *, peptides where a negative percent exchange was corrected to zero (<2% magnitude correction); **, peptides that were missing a replicate at the 30 min timepoint.

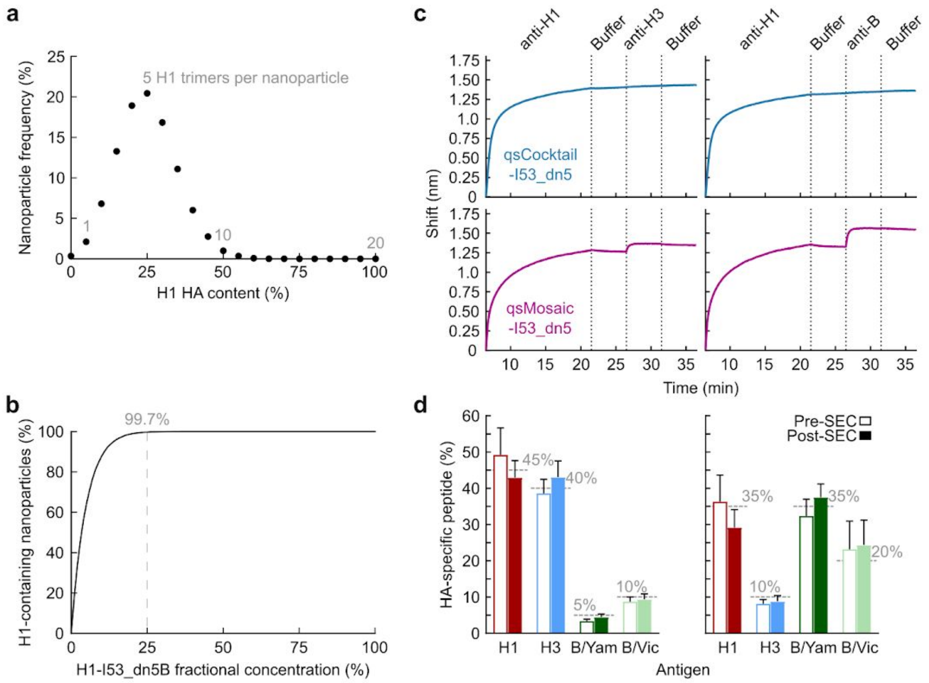


**Extended Data 3 | Controllable co-display of multiple antigenic variants on two-component nanoparticle immunogens. a,** Numerical approximation of the H1 HA content of individual qsMosaic-I53_dn5 nanoparticles assuming an equimolar quadrivalent *in vitro* assembly reaction (i.e., 25% of the input HA-I53_dn5B trimers bear H1 HA) and random incorporation of each HA-I53_dn5B trimer at each of the 20 trimeric positions into the nanoparticle. A distribution centered on 25% valency (5 H1 HA trimers per nanoparticle) is observed. **b,** Calculation of the fraction of individual mosaic nanoparticles displaying at least one H1 HA trimer as a function of the fractional concentration of H1-I53-dn5B in the *in vitro* assembly reaction ([H1]), expressed as: $1 - (1 - [H1])^{20}$. At the 25% fractional concentration used to assemble qsMosaic-I53_dn5, 99.7% of the individual nanoparticles are expected to display at least one H1 HA trimer. **c,** Sandwich BLI comparing qsCocktail-I53_dn5 and qsMosaic-I53_dn5. Biotinylated 5J8 immobilized on streptavidin probes was used to capture H1-containing nanoparticles from each sample. The captured particles were then exposed to antibodies specific to H3 (CR8020; left) or influenza B HA (CR8071; right). **d,** Quantitation of HA antigen content in two distinct qsMosaic-I53_dn5 nanoparticles with non-uniform antigen ratios before and after preparative SEC by peptide mass spectrometry. Dashed grey lines represent the fractional concentration of each HA in the *in vitro* assembly reactions used to prepare the mosaic nanoparticle immunogens, and error bars represent the standard deviation of measurements across four unique peptides from each HA. The peptides used to quantify each HA are provided in Supplementary Table 3.

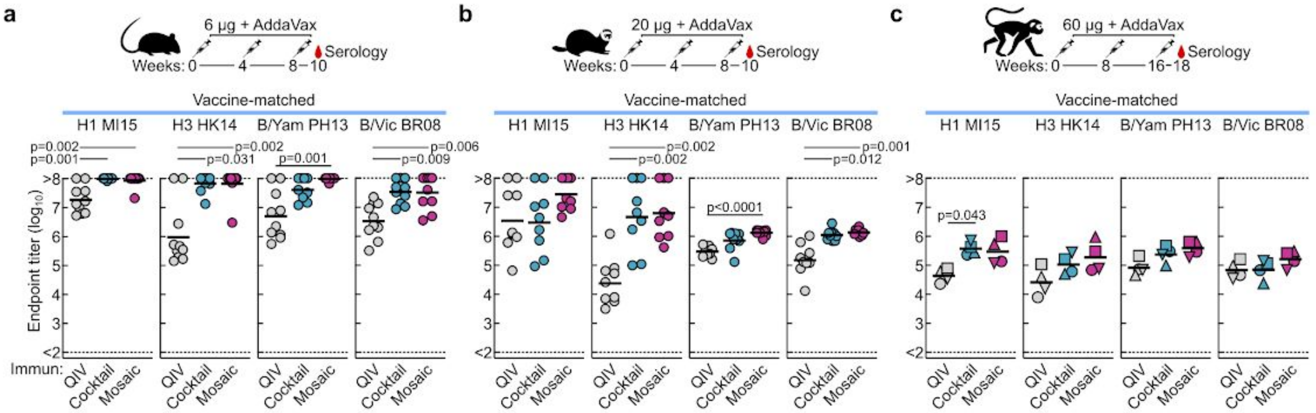


**Extended Data Fig. 4 | Vaccine-elicited HA-specific antibody titers against vaccine-matched antigens.** HA-specific antibody titers in immunized **a,** mice, **b,** ferrets, and **c,** NHPs. Immunization schemes are shown at the top of each panel. All immunizations were given intramuscularly with AddaVax. Groups of BALB/cJ mice (*N* = 10), Finch ferrets (*N* = 9), and rhesus macaques (*N* = 4) were used in each experiment. Antibody titers are expressed as endpoint dilutions. Each symbol represents an individual animal and the horizontal bar indicates the geometric mean of the group. Individual NHPs are identified by unique symbols. Statistical analysis was performed using nonparametric Kruskal–Wallis test with Dunn's multiple comparisons. All animal experiments except for NHP were performed at least twice and representative data are shown.

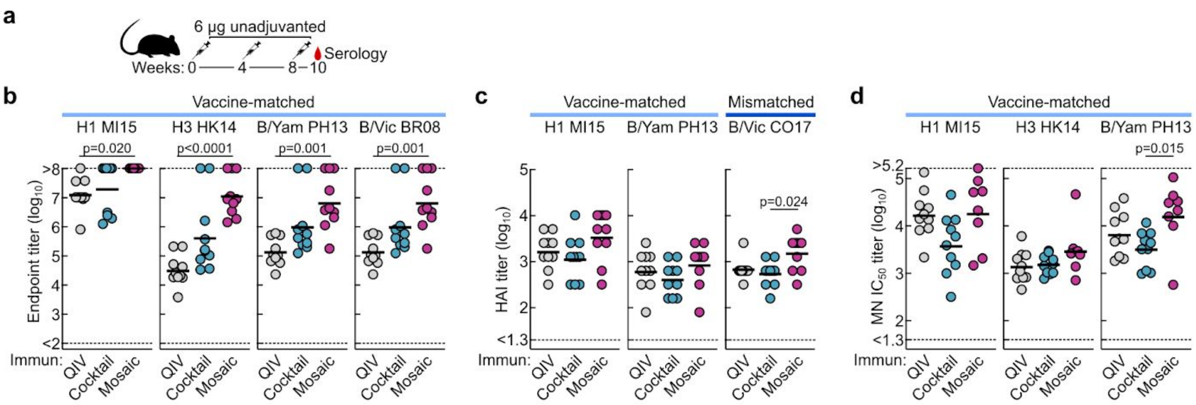


**Extended Data Fig. 5 | Antibody responses against vaccine-matched antigens and viruses elicited by unadjuvanted vaccines. a,** Immunization scheme. All immunizations were given intramuscularly without adjuvant. Groups of BALB/cJ mice (*N* = 10) were used. **b,** HA-specific antibody, **c,** hemagglutination inhibition (HAI), and **d,** microneutralization titers in immune sera. Microneutralization titers are reported as half maximal inhibitory dilution ($IC_{50}$). Each symbol represents an individual animal and the horizontal bar indicates the geometric mean of the group. Statistical analysis was performed using nonparametric Kruskal–Wallis test with Dunn's multiple comparisons. The experiment was performed twice and representative data are shown.

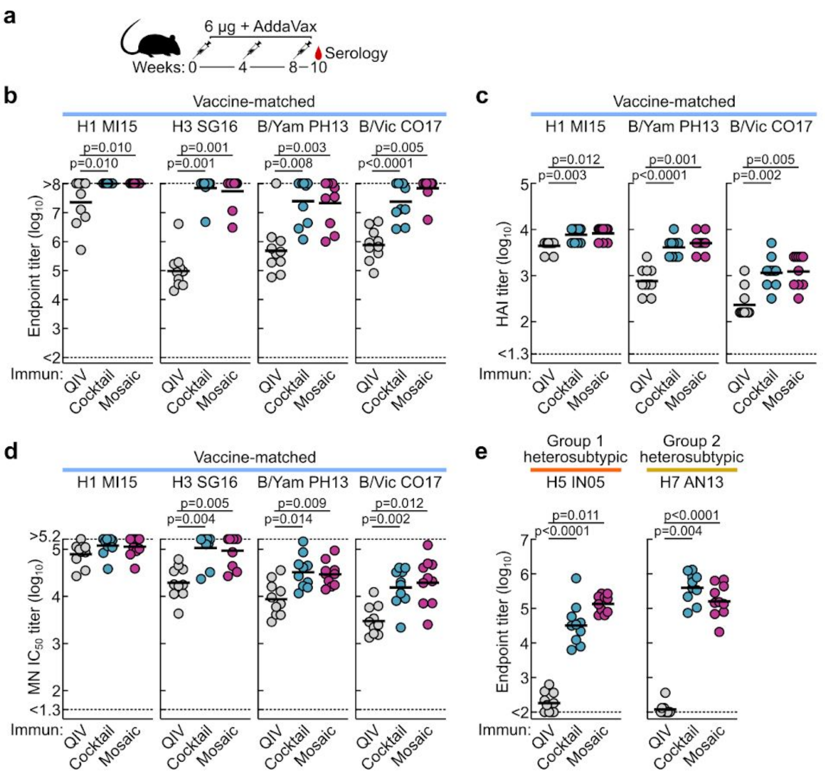


**Extended Data Fig. 6 | Antibody responses against vaccine-matched antigens and viruses elicited by 2018-2019 vaccines. a,** Immunization scheme. The commercial QIV, qsCocktail-I53_dn5, and qsMosaic-I53_dn5 vaccines used in this study comprised the WHO-recommended 2018-2019 vaccine strains. Sequences for the HA-I53_dn5B fusion proteins—H1-I53_dn5, SG16-I53_dn5 (updated H3), B/Yam-I53_dn5, and CO17-I53_dn5 (updated B/Vic)—are provided in Supplementary Table 1. All immunizations were given intramuscularly with AddaVax. Groups of BALB/cJ mice ($N$ = 10) were used. **b,** HA-specific antibody, **c,** hemagglutination inhibition (HAI), and **d,** microneutralization titers in immune sera. Microneutralization titers are reported as half maximal inhibitory dilution ($IC_{50}$). **e,** Heterosubtypic HA-specific antibody titers in immune sera. Each symbol represents an individual animal and the horizontal bar indicates the geometric mean of the group. Statistical analysis was performed using nonparametric Kruskal–Wallis test with Dunn's multiple comparisons. The animal experiment was performed once.

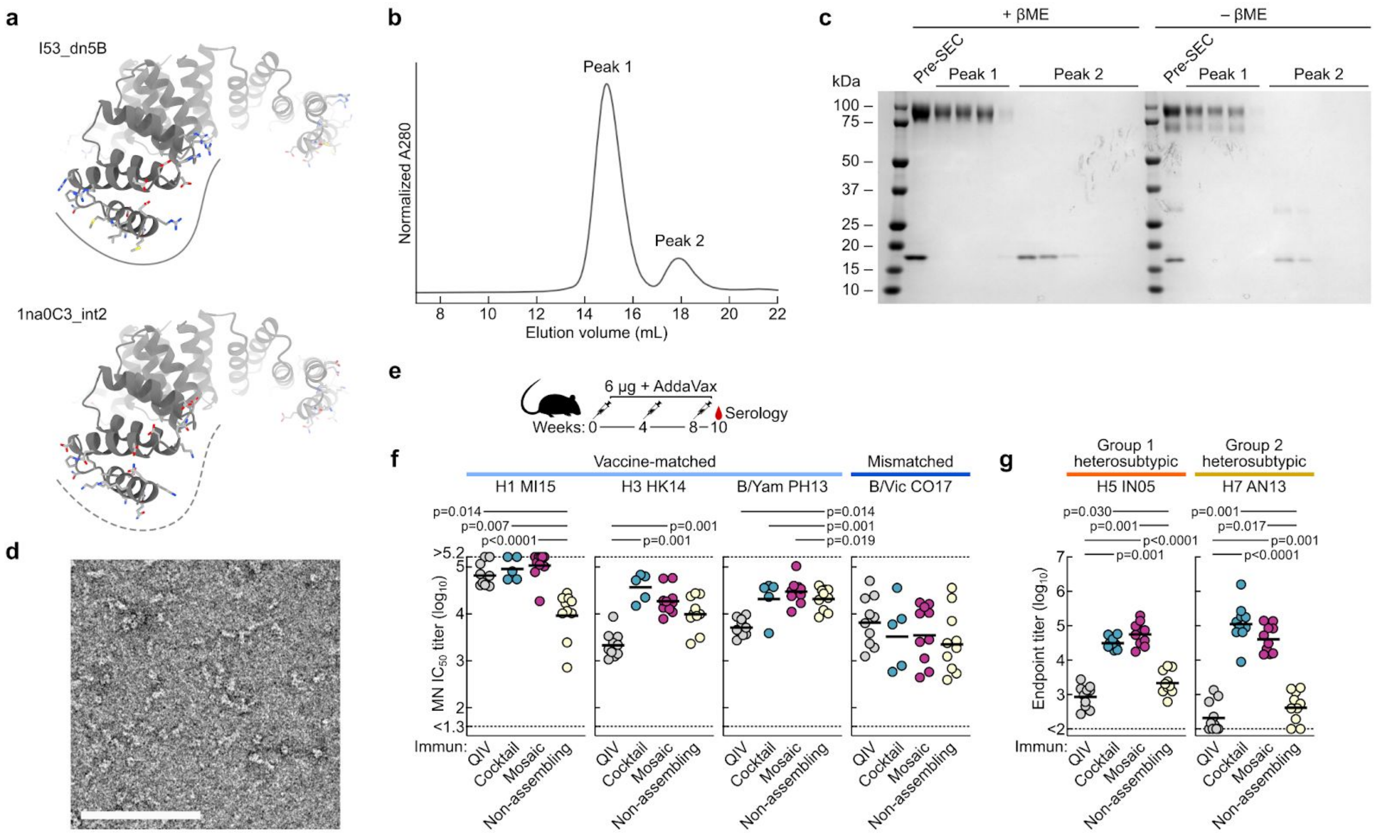


**Extended Data Fig. 7 | Antibody responses elicited by a non-assembling immunogen.** **a,** Model of the I53_dn5B trimer, with the computationally designed interface that drives nanoparticle assembly indicated by the solid line (top), and the 1na0C3_int2 trimer, in which the interface mutations were reverted to their original identities (bottom). The dotted line indicates the inability of this molecule to drive nanoparticle assembly. **b,** Analytical SEC of the non-assembling immunogen (a mixture of four HA-1na0C3_int2 trimers with pentameric I53_dn5A) using a Superose 6 Increase 10/300 GL column. Only unassembled oligomeric components were observed. **c,** Reducing and non-reducing SDS-PAGE analysis of the non-assembling immunogen before and after analytical SEC. **d,** Negative stain EM of the non-assembling immunogen, which confirmed the absence of higher-order structures indicated by analytical SEC. Scale bar, 100 nm. **e,** Immunization scheme in mice. All immunizations were given intramuscularly with AddaVax. Groups of BALB/cJ mice (*N* = 10) were used in the experiment. **f,** Microneutralization titers in immune sera against vaccine-matched or slightly mismatched viruses. Microneutralization titers are reported as half maximal inhibitory dilution ($IC_{50}$). **g,** Cross-reactive antibody titers in immune sera. Each symbol represents an individual animal and the horizontal bar indicates the geometric mean of the group. Statistical analysis was performed using nonparametric Kruskal–Wallis test with Dunn's multiple comparisons. The animal experiment was performed once.

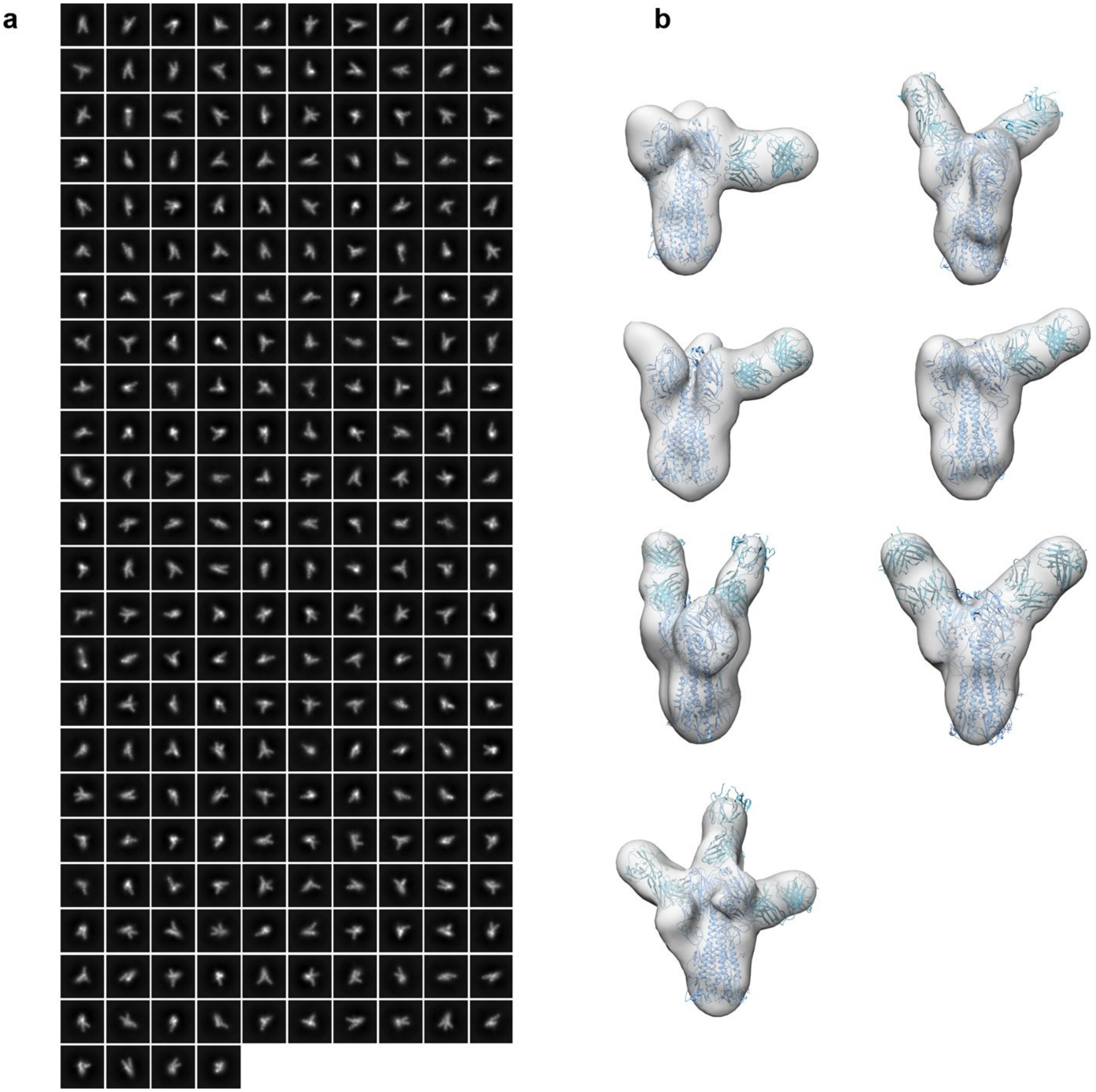


**Extended Data Fig. 8 | EM analysis of negatively stained vaccine-matched H1-foldon complexed with polyclonal Fabs elicited by qsCocktail-I53_dn5. a,** Representative reference-free 2D class averages. 4,112 micrographs were collected and 847,873 particles were used for 2D classification. The frequencies of HA/Fab complexes observed by cryo-EM containing Fab fragments bound to RBD (81%), VE (18%), or stem (1%) domains are presented as pie charts in Fig. 5f. Single complexes containing Fabs of multiple specificities were counted once against each specificity. **b,** Seven representative 3D reconstructions for immune complexes with Fabs are shown. The coordinates of an H1 HA crystal structure (PDB 1RUZ) and a Fab fragment (PDB 3GBN) were fitted into the EM densities. Light blue ribbons, H1 HA; cyan ribbons, polyclonal Fabs.

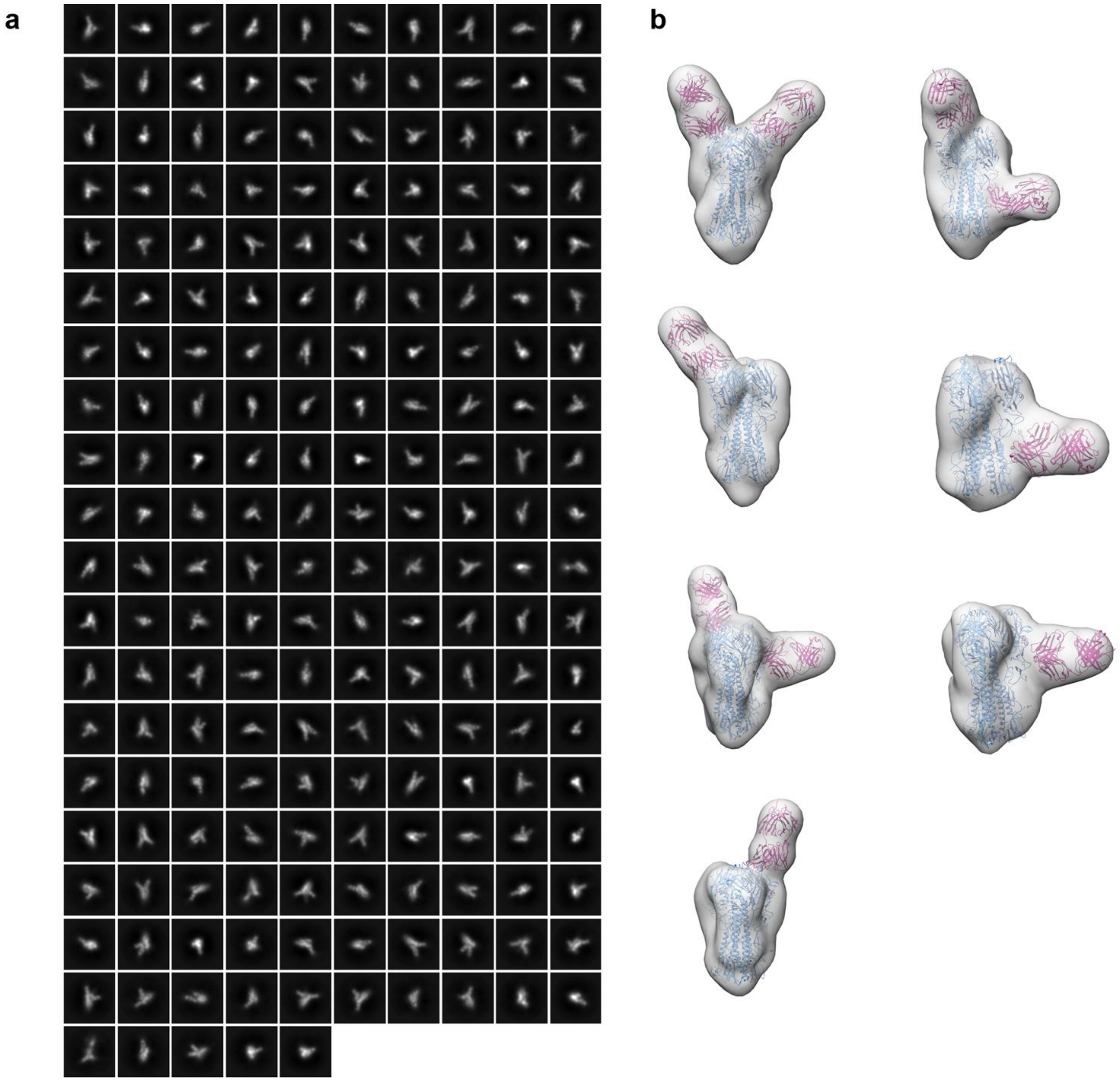


**Extended Data Fig. 9 | EM analysis of negatively stained vaccine-matched H1-foldon complexed with polyclonal Fabs elicited by qsMosaic-I53_dn5. a,** Representative reference-free 2D class averages. 3,237 micrographs were collected and 997,557 particles were used for 2D classification. The frequencies of HA/Fab complexes observed by cryo-EM containing Fab fragments bound to RBD (69%), VE (24%), or stem (7%) domains are presented as pie charts in Fig. 5f. Single complexes containing Fabs of multiple specificities were counted once against each specificity. **b,** Seven representative 3D reconstructions for immune complexes with Fabs are shown. The coordinates of an H1 HA crystal structure (PDB 1RUZ) and a Fab fragment (PDB 3GBN) were fitted into the EM densities. Light blue ribbons, H1 HA; violet ribbons, polyclonal Fabs.

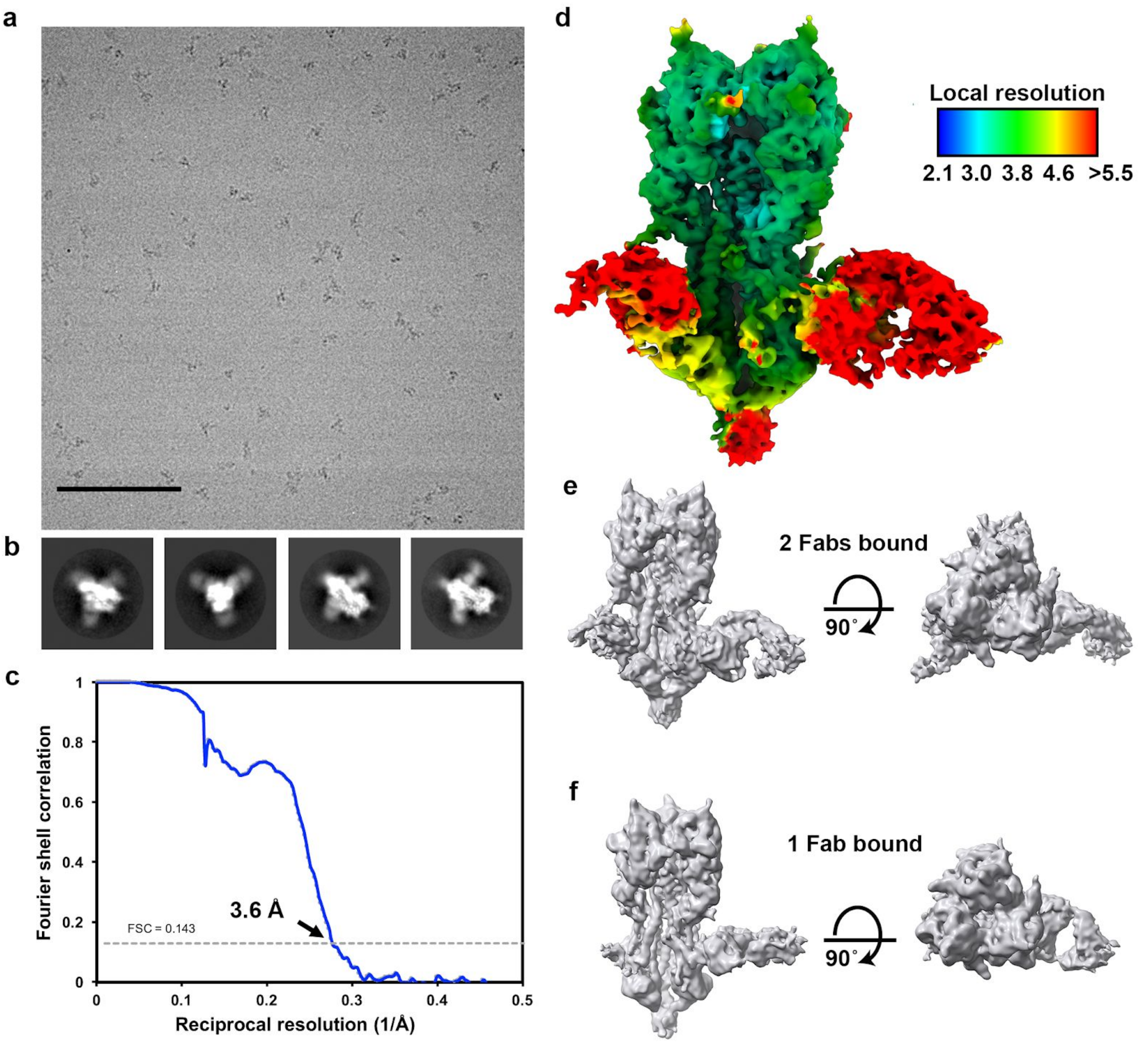


**Extended Data Fig. 10 | CryoEM analysis of heterosubtypic H5 HA-foldon in complex with polyclonal antibody Fab fragments elicited by qsMosaic-I53_dn5. a,** Representative cryo-electron micrograph. Scale bar, 100 nm. **b,** Reference-free 2D class averages. **c,** Gold-standard Fourier shell correlation curve for the asymmetric reconstruction shown in **d**. **d,** Asymmetric cryo-EM reconstruction of H5 HA-foldon with polyclonal antibody Fab fragments elicited by qsMosaic-I53_dn5 bound to all three HA subunits. The reconstruction is the same as that shown in Fig. 5e,g, but here is colored by local resolution. **e,** Two orthogonal orientations of an unsharpened asymmetric cryo-EM reconstruction of H5 HA-foldon with polyclonal antibody Fab fragments elicited by qsMosaic-I53_dn5 bound to two HA subunits at 4.1 Å resolution. **f,** Two orthogonal orientations of an unsharpened asymmetric cryo-EM reconstruction of H5 HA-foldon with polyclonal antibody Fab fragments elicited by qsMosaic-I53_dn5 bound to one HA subunit at 4.0 Å resolution. Note the nearly orthogonal angles of approach indicated by the density in panels **e** and **f**.
