## Supplementary Information for "Elicitation of broadly protective immunity to influenza by multivalent hemagglutinin nanoparticle vaccines"

**Supplementary Table 1 | Amino acid sequences for novel proteins used in this study.**

**Non-antigen-bearing nanoparticle components**

>I53_dn5A
MGKYDGSKLRIGILHARWNAEIILALVLGALKRLQEFGVKRENIIIETVPGSFELPYGSKLFVEKQKRLGKPLDAIIPIG
VLIKGSTMHFEYICDSTTHQLMKLNFELGIPVIFGVLTCLTDEQAEARAGLIEGKMHNHGEDWGAAAVEMATKFNLE
HHHHHH

>I53_dn5B
MEEAELAYLLGELAYKLGEYRIAIRAYRIALKRDPNNAEAWYNLGNAYYKQGRYREAIEYYQKALELDPNNAEAWY
NLGNAYYERGEYEEAIEYYRKALRLDPNNADAMQNLLNAKMREEGGWELQGSLEHHHHHH

**HA-I53_dn5B trimers**

>H1-I53_dn5B: A/Michigan/45/2015 HA 1-676 Y98F no lkr dn5B.SA.WELQ-H
MKAILVVLLYTFTTANADTLCIGYHANNSTDTVDTVLEKNVTVTHSVNLLEDKHNGKLCKLRGVAPLHLGKCNIAGWI
LGNPECESLSTASSWSYIVETSNSDNGTCFPGDFINYEELREQLSSVSSFERFEIFPKTSSWPNHDSNKGVTAACP
HAGAKSFYKNLIWLVKKGNSYPKLNQSYINDKGKEVLVLWGIHHPSTTADQQSLYQNADAYVFVGTSRYSKKFKPE
IATRPKVRDQEGRMNYYWTLVEPGDKITFEATGNLVVPRYAFTMERNAGSGIIISDTPVHDCNTTCQTPEGAINTSL
PFQNIHPITIGKCPKYVKSTKLRLATGLRNVPSIQSRGLFGAIAGFIEGGWTGMVDGWYGYHHQNEQGSGYAADLK
STQNAIDKITNKVNSVIEKMNTQFTAVGKEFNHLEKRIENLNKKVDDGFLDIWTYNAELLVLLENERTLDYHDSNVKN
LYEKVRNQLKNNAKEIGNGCFEFYHKCDNTCMESVKNGTYDYPKYSEEAKLNREKIDGVSAEEAELAYLLGELAYK
LGEYRIAIRAYRIALKRDPNNAEAWYNLGNAYYKQGRYREAIEYYQKALELDPNNAEAWYNLGNAYYERGEYEEAI
EYYRKALRLDPNNADAMQNLLNAKMREEGGWELQHHHHHH

>H3-I53_dn5B: A/Hong Kong/4801/2014 HA 1-676 Y98F GG lkr dn5B.SA.WELQ-H
MKTIIALSYILCLVFAQKIPGNDNSTATLCLGHHAVPNGTIVKTITNDRIEVTNATELVQNSSIGEICDSPHQILDGENC
TLIDALLGDPQCDGFQNKKWDLFVERSKAYSNCFPYDVPDYASLRSLVASSGTLEFNNESFNWTGVTQNGTSSAC
IRRSSSSFFSRLNWLTHLNYTYPALNVTMPNNEQFDKLYIWGVHHPGTDKDQIFLYAQSSGRITVSTKRSQQAVIPN
IGSRPRIRDIPSRISIYWTIVKPGDILLINSTGNLIAPRGYFKIRSGKSSIMRSDAPIGKCKSECITPNGSIPNDKPFQNV
NRITYGACPRYVKHSTLKLATGMRNVPEKQTRGIFGAIAGFIENGWEGMVDGWYGFRHQNSEGRGQAADLKSTQ
AAIDQINGKLNRLIGKTNEKFHQIEKEFSEVEGRIQDLEKYVEDTKIDLWSYNAELLVALENQHTIDLTDSEMNKLFEK
TKKQLRENAEDMGNGCFKIYHKCDNACIGSIRNGTYDHNVYRDEALNNRFQIKGVGGSAEEAELAYLLGELAYKLG
EYRIAIRAYRIALKRDPNNAEAWYNLGNAYYKQGRYREAIEYYQKALELDPNNAEAWYNLGNAYYERGEYEEAIEY
YRKALRLDPNNADAMQNLLNAKMREEGGWELQHHHHHH

>B/Yam-I53_dn5B: B/Phuket/3073/2013 HA 1-674 no lkr dn5B.SA.WELQ-H
MKAIIVLLMVVTSNADRICTGITSSNSPHVVKTATQGEVNVTGVIPLTTTPTKSYFANLKGTRTRGKLCPDCLNCTDL
DVALGRPMCVGTTPSAKASILHEVRPVTSGCFPIMHDRTKIRQLPNLLRGYEKIRLSTQNVIDAEKAPGGPYRLGTS
GSCPNATSKIGFFATMAWAVPKDNYKNATNPLTVEVPYICTEGEDQITVWGFHSDNKTQMKSLYGDSNPQKFTSS
ANGVTTHYVSQIGDFPDQTEDGGLPQSGRIVVDYMMQKPGKTGTIVYQRGVLLPQKVWCASGRSKVIKGSLPLIG
EADCLHEEYGGLNKSKPYYTGKHAKAIGNCPIWVKTPLKLANGTKYRPPAKLLKERGFFGAIAGFLEGGWEGMIAG
WHGYTSHGAHGVAVAADLKSTQEAINKITKNLNSLSELEVKNLQRLSGAMDELHNEILELDEKVDDLRADTISSQIEL
AVLLSNEGIINSEDEHLLALERKLKKMLGPSAVDIGNGCFETKHKCNQTCLDRIAAGTFNAGEFSLPTFDSLNITAAS
AEEAELAYLLGELAYKLGEYRIAIRAYRIALKRDPNNAEAWYNLGNAYYKQGRYREAIEYYQKALELDPNNAEAWY
NLGNAYYERGEYEEAIEYYRKALRLDPNNADAMQNLLNAKMREEGGWELQHHHHHH

>B/Vic-I53_dn5B: B/Brisbane/60/2008 HA 1-674 no lkr dn5B.SA.WELQ-H
MKAIIVLLMVVTSNADRICTGITSSNSPHVVKTATQGEVNVTGVIPLTTTPTKSHFANLKGTETRGKLCPKCLNCTDL
DVALGRPKCTGKIPSARVSILHEVRPVTSGCFPIMHDRTKIRQLPNLLRGYEHIRLSTHNVINAENAPGGPYKIGTSG
SCPNITNGNGFFATMAWAVPKNDKNKTATNPLTIEVPYICTEGEDQITVWGFHSDNETQMAKLYGDSKPQKFTSSA
NGVTTHYVSQIGGFPNQTEDGGLPQSGRIVVDYMVQKSGKTGTITYQRGILLPQKVWCASGRSKVIKGSLPLIGEA

DCLHEKYGGLNKSKPYYTGEHAKAIGNCPIWVKTPLKLANGTKYRPPAKLLKERGFFGAIAGFLEGGWEGMIAGW
HGYTSHGAHGVAVAADLKSTQEAINKITKNLNSLSELEVKNLQRLSGAMDELHNEILELDEKVDDLRADTISSQIELA
VLLSNEGIINSEDEHLLALERKLKKMLGPSAVEIGNGCFETKHKCNQTCLDRIAAGTFDAGEFSLPTFDSLNITAASA
EEAELAYLLGELAYKLGEYRIAIRAYRIALKRDPNNAEAWYNLGNAYYKQGRYREAIEYYQKALELDPNNAEAWYNL
GNAYYERGEYEEAIEYYRKALRLDPNNADAMQNLLNAKMREEGGWELQHHHHHH

>SI16-I53_dn5B: A/Singapore/INFIMH-16-0019/2016 HA 1-676 Y98F GG lkr dn5B.SA.WELQ-H
MKTIIALSYILCLVFAQKIPGNDNSTATLCLGHHAVPNGTIVKTITNDRIEVTNATELVQNSSIGEICDSPHQILDGENC
TLIDALLGDPQCDGFQNKKWDLFVERSKAYSNCFPYDVPDYASLRSLVASSGTLEFKNESFNWTGVTQNGTSSAC
IRGSSSSFFSRLNWLTHLNYTYPALNVTMPNKEQFDKLYIWGVHHPGTDKDQIFLYAQSSGRITVSTKRSQQAVIPN
IGSRPRIRDIPSRISIYWTIVKPGDILLINSTGNLIAPRGYFKIRSGKSSIMRSDAPIGKCKSECITPNGSIPNDKPFQNV
NRITYGACPRYVKHSTLKLATGMRNVPEKQTRGIFGAIAGFIENGWEGMVDGWYGFRHQNSEGRGQAADLKSTQ
AAIDQINGKLNRLIGKTNEKFHQIEKEFSEVEGRVQDLEKYVEDTKIDLWSYNAELLVALENQHTIDLTDSEMNKLFE
KTKKQLRENAEDMGNGCFKIYHKCDNACIESIRNETYDHNVYRDEALNNRFQIKGVGGSAEEAELAYLLGELAYKL
GEYRIAIRAYRIALKRDPNNAEAWYNLGNAYYKQGRYREAIEYYQKALELDPNNAEAWYNLGNAYYERGEYEEAIE
YYRKALRLDPNNADAMQNLLNAKMREEGGWELQHHHHHH

>CO17-I53_dn5B: B/Colorado/06/2017 HA 1-672 no lkr dn5B.SA.WELQ-H
MKAIIVLLMVVTSSADRICTGITSSNSPHVVKTATQGEVNVTGVIPLTTTPTKSHFANLKGTETRGKLCPKCLNCTDL
DVALGRPKCTGKIPSARVSILHEVRPVTSGCFPIMHDRTKIRQLPNLLRGYEHVRLSTHNVINAEGAPGGPYKIGTS
GSCPNITNGNGFFATMAWAVPDKNKTATNPLTIEVPYVCTEGEDQITVWGFHSDNETQMAKLYGDSKPQKFTSSA
NGVTTHYVSQIGGFPNQTEDGGLPQSGRIVVDYMVQKSGKTGTITYQRGILLPQKVWCASGRSKVIKGSLPLIGEA
DCLHEKYGGLNKSKPYYTGEHAKAIGNCPIWVKTPLKLANGTKYRPPAKLLKERGFFGAIAGFLEGGWEGMIAGW
HGYTSHGAHGVAVAADLKSTQEAINKITKNLNSLSELEVKNLQRLSGAMDELHNEILELDEKVDDLRADTISSQIELA
VLLSNEGIINSEDEHLLALERKLKKMLGPSAVEIGNGCFETKHKCNQTCLDKIAAGTFDAGEFSLPTFDSLNITAASA
EEAELAYLLGELAYKLGEYRIAIRAYRIALKRDPNNAEAWYNLGNAYYKQGRYREAIEYYQKALELDPNNAEAWYNL
GNAYYERGEYEEAIEYYRKALRLDPNNADAMQNLLNAKMREEGGWELQHHHHHH

**HA-1na0C3_int2 (non-assembling) trimers**

>H1-1na0C3_int2: A/Michigan/45/2015 HA 1-676 Y98F no lkr 1na0C3_int2.SA.WELQ-H
MKAILVVLLYTFTTANADTLCIGYHANNSTDTVDTVLEKNVTVTHSVNLLEDKHNGKLCKLRGVAPLHLGKCNIAGWI
LGNPECESLSTASSWSYIVETSNSDNGTCFPGDFINYEELREQLSSVSSFERFEIFPKTSSWPNHDSNKGVTAACP
HAGAKSFYKNLIWLVKKGNSYPKLNQSYINDKGKEVLVLWGIHHPSTTADQQSLYQNADAYVFVGTSRYSKKFKPE
IATRPKVRDQEGRMNYYWTLVEPGDKITFEATGNLVVPRYAFTMERNAGSGIIISDTPVHDCNTTCQTPEGAINTSL
PFQNIHPITIGKCPKYVKSTKLRLATGLRNVPSIQSRGLFGAIAGFIEGGWTGMVDGWYGYHHQNEQGSGYAADLK
STQNAIDKITNKVNSVIEKMNTQFTAVGKEFNHLEKRIENLNKKVDDGFLDIWTYNAELLVLLENERTLDYHDSNVKN
LYEKVRNQLKNNAKEIGNGCFEFYHKCDNTCMESVKNGTYDYPKYSEEAKLNREKIDGVSAEEAELAYLLGELAYK
LGEYRIAIRAYRIALKRDPNNAEAWYNLGNAYYKQGDYDEAIEYYQKALELDPNNAEAWYNLGNAYYKQGDYDEAI
EYYQKALELDPNNAEAKQNLGNAKQKQGGGWELQHHHHHH

>H3-1na0C3_int2: A/Hong Kong/4801/2014 HA 1-676 Y98F GG lkr 1na0C3_int2.SA.WELQ-H
MKTIIALSYILCLVFAQKIPGNDNSTATLCLGHHAVPNGTIVKTITNDRIEVTNATELVQNSSIGEICDSPHQILDGENC
TLIDALLGDPQCDGFQNKKWDLFVERSKAYSNCFPYDVPDYASLRSLVASSGTLEFNNESFNWTGVTQNGTSSAC
IRRSSSSFFSRLNWLTHLNYTYPALNVTMPNNEQFDKLYIWGVHHPGTDKDQIFLYAQSSGRITVSTKRSQQAVIPN
IGSRPRIRDIPSRISIYWTIVKPGDILLINSTGNLIAPRGYFKIRSGKSSIMRSDAPIGKCKSECITPNGSIPNDKPFQNV
NRITYGACPRYVKHSTLKLATGMRNVPEKQTRGIFGAIAGFIENGWEGMVDGWYGFRHQNSEGRGQAADLKSTQ
AAIDQINGKLNRLIGKTNEKFHQIEKEFSEVEGRIQDLEKYVEDTKIDLWSYNAELLVALENQHTIDLTDSEMNKLFEK
TKKQLRENAEDMGNGCFKIYHKCDNACIGSIRNGTYDHNVYRDEALNNRFQIKGVGGSAEEAELAYLLGELAYKLG
EYRIAIRAYRIALKRDPNNAEAWYNLGNAYYKQGDYDEAIEYYQKALELDPNNAEAWYNLGNAYYKQGDYDEAIEY
YQKALELDPNNAEAKQNLGNAKQKQGGGWELQHHHHHH

>B/Yam-1na0C3_int2: B/Phuket/3073/2013 HA 1-674 no lkr 1na0C3_int2.SA.WELQ-H
MKAIIVLLMVVTSNADRICTGITSSNSPHVVKTATQGEVNVTGVIPLTTTPTKSYFANLKGTRTRGKLCPDCLNCTDL
DVALGRPMCVGTTPSAKASILHEVRPVTSGCFPIMHDRTKIRQLPNLLRGYEKIRLSTQNVIDAEKAPGGPYRLGTS
GSCPNATSKIGFFATMAWAVPKDNYKNATNPLTVEVPYICTEGEDQITVWGFHSDNKTQMKSLYGDSNPQKFTSS
ANGVTTHYVSQIGDFPDQTEDGGLPQSGRIVVDYMMQKPGKTGTIVYQRGVLLPQKVWCASGRSKVIKGSLPLIG
EADCLHEEYGGLNKSKPYYTGKHAKAIGNCPIWVKTPLKLANGTKYRPPAKLLKERGFFGAIAGFLEGGWEGMIAG
WHGYTSHGAHGVAVAADLKSTQEAINKITKNLNSLSELEVKNLQRLSGAMDELHNEILELDEKVDDLRADTISSQIEL
AVLLSNEGIINSEDEHLLALERKLKKMLGPSAVDIGNGCFETKHKCNQTCLDRIAAGTFNAGEFSLPTFDSLNITAAS
AEEAELAYLLGELAYKLGEYRIAIRAYRIALKRDPNNAEAWYNLGNAYYKQGDYDEAIEYYQKALELDPNNAEAWY
NLGNAYYKQGDYDEAIEYYQKALELDPNNAEAKQNLGNAKQKQGGGWELQHHHHHH

>B/Vic-1na0C3_int2: B/Brisbane/60/2008 HA 1-674 no lkr 1na0C3_int2.SA.WELQ-H
MKAIIVLLMVVTSNADRICTGITSSNSPHVVKTATQGEVNVTGVIPLTTTPTKSHFANLKGTETRGKLCPKCLNCTDL
DVALGRPKCTGKIPSARVSILHEVRPVTSGCFPIMHDRTKIRQLPNLLRGYEHIRLSTHNVINAENAPGGPYKIGTSG
SCPNITNGNGFFATMAWAVPKNDKNKTATNPLTIEVPYICTEGEDQITVWGFHSDNETQMAKLYGDSKPQKFTSSA
NGVTTHYVSQIGGFPNQTEDGGLPQSGRIVVDYMVQKSGKTGTITYQRGILLPQKVWCASGRSKVIKGSLPLIGEA
DCLHEKYGGLNKSKPYYTGEHAKAIGNCPIWVKTPLKLANGTKYRPPAKLLKERGFFGAIAGFLEGGWEGMIAGW
HGYTSHGAHGVAVAADLKSTQEAINKITKNLNSLSELEVKNLQRLSGAMDELHNEILELDEKVDDLRADTISSQIELA
VLLSNEGIINSEDEHLLALERKLKKMLGPSAVEIGNGCFETKHKCNQTCLDRIAAGTFDAGEFSLPTFDSLNITAASA
EEAELAYLLGELAYKLGEYRIAIRAYRIALKRDPNNAEAWYNLGNAYYKQGDYDEAIEYYQKALELDPNNAEAWYNL
GNAYYKQGDYDEAIEYYQKALELDPNNAEAKQNLGNAKQKQGGGWELQHHHHHH

**HA-foldon trimers (ELISA antigens and EMPEM)**

>H1 MI15: A/Michigan/45/2015 HA 1-676 Y98F FAH
MKAILVVLLYTFTTANADTLCIGYHANNSTDTVDTVLEKNVTVTHSVNLLEDKHNGKLCKLRGVAPLHLGKCNIAGWI
LGNPECESLSTASSWSYIVETSNSDNGTCFPGDFINYEELREQLSSVSSFERFEIFPKTSSWPNHDSNKGVTAACP
HAGAKSFYKNLIWLVKKGNSYPKLNQSYINDKGKEVLVLWGIHHPSTTADQQSLYQNADAYVFVGTSRYSKKFKPE
IATRPKVRDQEGRMNYYWTLVEPGDKITFEATGNLVVPRYAFTMERNAGSGIIISDTPVHDCNTTCQTPEGAINTSL
PFQNIHPITIGKCPKYVKSTKLRLATGLRNVPSIQSRGLFGAIAGFIEGGWTGMVDGWYGYHHQNEQGSGYAADLK
STQNAIDKITNKVNSVIEKMNTQFTAVGKEFNHLEKRIENLNKKVDDGFLDIWTYNAELLVLLENERTLDYHDSNVKN
LYEKVRNQLKNNAKEIGNGCFEFYHKCDNTCMESVKNGTYDYPKYSEEAKLNREKIDGVGSGYIPEAPRDGQAYV
RKDGEWVLLSTFLGSGLNDIFEAQKIEWHEGHHHHHH

>H3 HK14: A/Hong Kong/4801/2014 HA 1-676 Y98F FAH
MKTIIALSYILCLVFAQKIPGNDNSTATLCLGHHAVPNGTIVKTITNDRIEVTNATELVQNSSIGEICDSPHQILDGENC
TLIDALLGDPQCDGFQNKKWDLFVERSKAYSNCFPYDVPDYASLRSLVASSGTLEFNNESFNWTGVTQNGTSSAC
IRRSSSSFFSRLNWLTHLNYTYPALNVTMPNNEQFDKLYIWGVHHPGTDKDQIFLYAQSSGRITVSTKRSQQAVIPN
IGSRPRIRDIPSRISIYWTIVKPGDILLINSTGNLIAPRGYFKIRSGKSSIMRSDAPIGKCKSECITPNGSIPNDKPFQNV
NRITYGACPRYVKHSTLKLATGMRNVPEKQTRGIFGAIAGFIENGWEGMVDGWYGFRHQNSEGRGQAADLKSTQ
AAIDQINGKLNRLIGKTNEKFHQIEKEFSEVEGRIQDLEKYVEDTKIDLWSYNAELLVALENQHTIDLTDSEMNKLFEK
TKKQLRENAEDMGNGCFKIYHKCDNACIGSIRNGTYDHNVYRDEALNNRFQIKGVGSGYIPEAPRDGQAYVRKDG
EWVLLSTFLGSGLNDIFEAQKIEWHEGHHHHHH

>B/Yam PH13: B/Phuket/3073/2013 HA 1-676 FAH
MKAIIVLLMVVTSNADRICTGITSSNSPHVVKTATQGEVNVTGVIPLTTTPTKSYFANLKGTRTRGKLCPDCLNCTDL
DVALGRPMCVGTTPSAKASILHEVRPVTSGCFPIMHDRTKIRQLPNLLRGYEKIRLSTQNVIDAEKAPGGPYRLGTS
GSCPNATSKIGFFATMAWAVPKDNYKNATNPLTVEVPYICTEGEDQITVWGFHSDNKTQMKSLYGDSNPQKFTSS
ANGVTTHYVSQIGDFPDQTEDGGLPQSGRIVVDYMMQKPGKTGTIVYQRGVLLPQKVWCASGRSKVIKGSLPLIG
EADCLHEEYGGLNKSKPYYTGKHAKAIGNCPIWVKTPLKLANGTKYRPPAKLLKERGFFGAIAGFLEGGWEGMIAG
WHGYTSHGAHGVAVAADLKSTQEAINKITKNLNSLSELEVKNLQRLSGAMDELHNEILELDEKVDDLRADTISSQIEL
AVLLSNEGIINSEDEHLLALERKLKKMLGPSAVDIGNGCFETKHKCNQTCLDRIAAGTFNAGEFSLPTFDSLNITAAS
LGSGYIPEAPRDGQAYVRKDGEWVLLSTFLGSGLNDIFEAQKIEWHEGHHHHHH

>B/Vic BR08: B/Brisbane/60/2008 HA 1-676 FAH
MKAIIVLLMVVTSNADRICTGITSSNSPHVVKTATQGEVNVTGVIPLTTTPTKSHFANLKGTETRGKLCPKCLNCTDL
DVALGRPKCTGKIPSARVSILHEVRPVTSGCFPIMHDRTKIRQLPNLLRGYEHIRLSTHNVINAENAPGGPYKIGTSG
SCPNITNGNGFFATMAWAVPKNDKNKTATNPLTIEVPYICTEGEDQITVWGFHSDNETQMAKLYGDSKPQKFTSSA
NGVTTHYVSQIGGFPNQTEDGGLPQSGRIVVDYMVQKSGKTGTITYQRGILLPQKVWCASGRSKVIKGSLPLIGEA
DCLHEKYGGLNKSKPYYTGEHAKAIGNCPIWVKTPLKLANGTKYRPPAKLLKERGFFGAIAGFLEGGWEGMIAGW
HGYTSHGAHGVAVAADLKSTQEAINKITKNLNSLSELEVKNLQRLSGAMDELHNEILELDEKVDDLRADTISSQIELA
VLLSNEGIINSEDEHLLALERKLKKMLGPSAVEIGNGCFETKHKCNQTCLDRIAAGTFDAGEFSLPTFDSLNITAASL
GSGYIPEAPRDGQAYVRKDGEWVLLSTFLGSGLNDIFEAQKIEWHEGHHHHHH

>H5 IN05: A/Indonesia/05/2005 HA 1-676 Y98F delF FAH
MEKIVLLLAIVSLVKSDQICIGYHANNSTEQVDTIMEKNVTVTHAQDILEKTHNGKLCDLDGVKPLILRDCSVAGWLL
GNPMCDEFINVPEWSYIVEKANPTNDLCFPGSFNDYEELKHLLSRINHFEKIQIIPKSSWSDHEASSGVSSACPYLG
SPSFFRNVVWLIKKNSTYPTIKKSYNNTNQEDLLVLWGIHHPNDAAEQTRLYQNPTTYISIGTSTLNQRLVPKIATRS
KVNGQSGRMEFFWTILKPNDAINFESNGNFIAPEYAYKIVKKGDSAIMKSELEYGNCNTKCQTPMGAINSSMPFHNI
HPLTIGECPKYVKSNRLVLATGLRNSPQRESRGLFGAIAGFIEGGWQGMVDGWYGYHHSNEQGSGYAADKESTQ
KAIDGVTNKVNSIIDKMNTQFEAVGREFNNLERRIENLNKKMEDGFLDVWTYNAELLVLMENERTLDFHDSNVKNL
YDKVRLQLRDNAKELGNGCFEFYHKCDNECMESIRNGTYNYPQYSEEARLKREEISGVGSGYIPEAPRDGQAYVR
KDGEWVLLSTFLGSGLNDIFEAQKIEWHEGHHHHHH

>H6 TW13: A/Taiwan/2/2013 HA 1-676 Y98F FAH
MIAIVVIAILASAGKSDKICIGYHANNSTTQVDTLLEKNVTVTHSVELLENQKEKRFCKIMNKAPLDLKDCTIEGWILGN
PKCDLLLGDQSWSYIVERPNAQNGICFPGVLNELEELKAFIGSGERVERFEMFPKSTWAGVDTSRGVTNACPSYTI
DSSFYRNLVWIVKTDSATYPVIKGTYNNTGTQPILYFWGVHHPLDTTVQDNLYGSGDKYVRMGTESMNFAKSPEIA
ARPAVNGQRSRIDYYWSVLRPGETLNVESNGNLIAPWYAYKFVSTNKKGAVFKSDLPIENCDATCQTITGVLRTNK
TFQNVSPLWIGECPKYVKSESLRLATGLRNVPQIATRGIFGAIAGFIEGGWTGMIDGWYGYHHENSQGSGYAADR
ESTQKAIDGITNKVNSIINKMNTQFEAVDHEFSNLERRIGNLNKRMEDGFLDVWTYNAELLVLLENERTLDLHDANV
KNLYEKVKSQLRDNANDLGNGCFEFWHKCDNECMESVKNGTYDYPKYQKESKLNRQGIESVGSGYIPEAPRDGQ
AYVRKDGEWVLLSTFLGSGLNDIFEAQKIEWHEGHHHHHH

>H7 AN13: A/Anhui/1/2013 HA 1-676 Y98F FAH
MNTQILVFALIAIIPTNADKICLGHHAVSNGTKVNTLTERGVEVVNATETVERTNIPRICSKGKRTVDLGQCGLLGTIT
GPPQCDQFLEFSADLIIERREGSDVCFPGKFVNEEALRQILRESGGIDKEAMGFTYSGIRTNGATSACRRSGSSFY
AEMKWLLSNTDNAAFPQMTKSYKNTRKSPALIVWGIHHSVSTAEQTKLYGSGNKLVTVGSSNYQQSFVPSPGARP
QVNGLSGRIDFHWLMLNPNDTVTFSFNGAFIAPDRASFLRGKSMGIQSGVQVDANCEGDCYHSGGTIISNLPFQNI
DSRAVGKCPRYVKQRSLLLATGMKNVPEIPKGRGLFGAIAGFIENGWEGLIDGWYGFRHQNAQGEGTAADYKSTQ
SAIDQITGKLNRLIEKTNQQFELIDNEFNEVEKQIGNVINWTRDSITEVWSYNAELLVAMENQHTIDLADSEMDKLYE
RVKRQLRENAEEDGTGCFEIFHKCDDDCMASIRNNTYDHSKYREEAMQNRIQIDPVGSGYIPEAPRDGQAYVRKD
GEWVLLSTFLGSGLNDIFEAQKIEWHEGHHHHHH

>H10 JD13: A/Jiangxi Donghu/346/2013 HA 1-676 Y98F FAH
MYKIVVIIALLGAVKGLDKICLGHHAVANGTIVKTLTNEQEEVTNATETVESTGINRLCMKGRKHKDLGNCHPIGMLIG
TPACDLHLTGMWDTLIERENAIAYCFPGATVNVEALRQKIMESGGINKISTGFTYGSSINSAGTTRACMRNGGNSFY
AELKWLVSKSKGQNFPQTTNTYRNTDTAEHLIMWGIHHPSSTQEKNDLYGTQSLSISVGSSTYRNNFVPVVGARP
QVNGQSGRIDFHWTLVQPGDNITFSHNGGLIAPSRVSKLIGRGLGIQSDAPIDNNCESKCFWRGGSINTRLPFQNL
SPRTVGQCPKYVNRRSLMLATGMRNVPELIQGRGLFGAIAGFLENGWEGMVDGWYGFRHQNAQGTGQAADYKS
TQAAIDQITGKLNRLVEKTNTEFESIESEFSEIEHQIGNVINWTKDSITDIWTYQAELLVAMENQHTIDMADSEMLNLY
ERVRKQLRQNAEEDGKGCFEIYHACDDSCMESIRNNTYDHSQYREEALLNRLNINPVGSGYIPEAPRDGQAYVRK
DGEWVLLSTFLGSGLNDIFEAQKIEWHEGHHHHHH

>H1 stem: A/California/04/2009 stabilized stem FAH
MKAILVVLLYTFATANADTLCIGYHANNSTDTVDTVLEKNVTVTHSVNLGSGLRLATGLRNIPQRETRGLFGAIAGFI
EGGWTGMVDGWYGYHHQNEQGSGYAADLKSTQNAIDEITNMVNSVIEKMGSGGSGTDLAELLVLLLNQWTLLYH

DSNVKNLYEKVRSQLKNNAKEIGNGCFEFYHKCDNTCMESVKNGTYDYPKYSEEAKLNREEIDGSGYIPEAPRDG
QAYVRKDGEWVLLSTFLGSGLNDIFEAQKIEWHEGHHHHHH

>H3 stem: A/Finland/486/2004 stabilized stem FAH
MKTIIALSYILCLVFAQKLPGNDNSTATLCLGHHAVPNGTIVKTITNDQIEVTNATELVFPGCGVLKLATGMRNVPEK
QTRGIFGAIAGFIENGWEGMVDGWYGFRHQNSEGIGQAADLKSTQAAINQINGMVNRVIALMAQGGPDCYLAELL
VALLNQHVIDLTDSEMRKLFERTKKQLRENAEDMGNGCFKIYHKCDNACIGSIRNGTYDHDVYRDEALNNRFQIKG
GPGSGYIPEAPRDGQAYVRKDGEWVLLSTFLGSGLNDIFEAQKIEWHEGHHHHHH

>H3 SG16: A/Singapore/INFIMH-16-0019/2016 HA 1-676 Y98F FAH
MKTIIALSYILCLVFAQKIPGNDNSTATLCLGHHAVPNGTIVKTITNDRIEVTNATELVQNSSIGEICDSPHQILDGENC
TLIDALLGDPQCDGFQNKKWDLFVERSKAYSNCFPYDVPDYASLRSLVASSGTLEFKNESFNWTGVTQNGTSSAC
IRGSSSSFFSRLNWLTHLNYTYPALNVTMPNKEQFDKLYIWGVHHPGTDKDQIFLYAQSSGRITVSTKRSQQAVIPN
IGSRPRIRDIPSRISIYWTIVKPGDILLINSTGNLIAPRGYFKIRSGKSSIMRSDAPIGKCKSECITPNGSIPNDKPFQNV
NRITYGACPRYVKHSTLKLATGMRNVPEKQTRGIFGAIAGFIENGWEGMVDGWYGFRHQNSEGRGQAADLKSTQ
AAIDQINGKLNRLIGKTNEKFHQIEKEFSEVEGRVQDLEKYVEDTKIDLWSYNAELLVALENQHTIDLTDSEMNKLFE
KTKKQLRENAEDMGNGCFKIYHKCDNACIESIRNETYDHNVYRDEALNNRFQIKGVGSGYIPEAPRDGQAYVRKD
GEWVLLSTFLGSGLNDIFEAQKIEWHEGHHHHHH

>B/Vic CO17: B/Colorado/06/2017 HA 1-676 FAH
MKAIIVLLMVVTSSADRICTGITSSNSPHVVKTATQGEVNVTGVIPLTTTPTKSHFANLKGTETRGKLCPKCLNCTDL
DVALGRPKCTGKIPSARVSILHEVRPVTSGCFPIMHDRTKIRQLPNLLRGYEHVRLSTHNVINAEGAPGGPYKIGTS
GSCPNITNGNGFFATMAWAVPDKNKTATNPLTIEVPYVCTEGEDQITVWGFHSDNETQMAKLYGDSKPQKFTSSA
NGVTTHYVSQIGGFPNQTEDGGLPQSGRIVVDYMVQKSGKTGTITYQRGILLPQKVWCASGRSKVIKGSLPLIGEA
DCLHEKYGGLNKSKPYYTGEHAKAIGNCPIWVKTPLKLANGTKYRPPAKLLKERGFFGAIAGFLEGGWEGMIAGW
HGYTSHGAHGVAVAADLKSTQEAINKITKNLNSLSELEVKNLQRLSGAMDELHNEILELDEKVDDLRADTISSQIELA
VLLSNEGIINSEDEHLLALERKLKKMLGPSAVEIGNGCFETKHKCNQTCLDKIAAGTFDAGEFSLPTFDSLNITAASL
GSGYIPEAPRDGQAYVRKDGEWVLLSTFLGSGLNDIFEAQKIEWHEGHHHHHH

Appended sequences including the foldon trimerization domain, WELQut protease recognition sites, Avi tags, and hexa-histidine tags are underlined.

**Supplementary Table 2 | Viruses used in HAI, microneutralization, and challenge experiments.**

| Subtype | Virus | Short name | Type of experiment | Figure |
|---|---|---|---|---|
| H1N1 | A/Michigan/45/2015 | MI15; 2015 | HAI; Microneutralization | 2c,d; 3c,d; 5b |
| | A/Boston/YGA-01050/2012 | 2012 | Microneutralization | 3c,d |
| | A/California/07/2009 | 2009 | Microneutralization | 3c,d |
| | A/Weiss/1943 | 1943 | Microneutralization | 3c,d |
| | A/Malaysia/1954 | 1954 | Microneutralization | 3c,d |
| | A/USSR/90/1977 | 1977 | Microneutralization | 3c,d |
| | A/Memphis/4/1987 | 1987 | Microneutralization | 3c,d |
| | A/New York/146/2000 | 2000 | Microneutralization | 3c,d |
| | A/New Caledonia/20/1999 | 1999 | Microneutralization | 3c,d |
| | A/Solomon Islands/03/2006 | 2006 | Microneutralization | 3c,d |
| | A/Puerto Rico/8/1934 | PR8 | Challenge (mice) | 3f |
| H3N2 | A/Hong Kong/4801/2014 | HK14; 2014 | Microneutralization | 2d; 3c,e |
| | A/Switzerland/9715293/2013 | 2013 | Microneutralization | 3c,e |
| | A/Texas/50/2012 | 2012 | Microneutralization | 3c,e |
| | A/Victoria/361/2011 | 2011 | Microneutralization | 3c,e |
| | A/Brisbane/10/2007 | 2007 | Microneutralization | 3c,e |
| | A/Wisconsin/67/2005 | 2005 | Microneutralization | 3c,e |
| | A/Fujian/411/2002 | 2002 | Microneutralization | 3c,e |
| | A/Moscow/10/1999 | 1999 | Microneutralization | 3c,e |
| | A/Shangdong/9/1993 | 1993 | Microneutralization | 3c,e |
| | A/Philippines/2/1982 | PH82 | Challenge (mice) | 3g |
| | A/Singapore/INFIMH-16-0019/2016 | SG16 | Microneutralization | ED 6 |
| H5N1 | A/Vietnam/1203/2004 | VN04 | Microneutralization; Challenge (mice, ferrets) | 4d-f |
| H7N9 | A/Anhui/1/2013 | AN13 | Challenge (mice, ferrets) | 4d-f |
| B/Yam | B/Phuket/3073/2013 | PH13 | HAI; Microneutralization | 2c,d |
| B/Vic | B/Colorado/06/2017 | CO17 | HAI; Microneutralization | 2c, ED 5-7 |

**Supplementary Table 3 | Peptides used in label-free quantitation of HA content in mosaic nanoparticles and data for post-SEC quantitation of HA content in qsMosaic-I53_dn5 (related to Fig. 1g).**

| HA construct | Unique peptide sequence[a] | Standard equimolar mix (area) | qsMosaic-I53_dn5 post-SEC (area) | Relative ratio (qsMosaic-I53_dn5 /standard) | Average relative ratio and standard deviation | qsMosaic-I53_dn5 post-SEC composition[b] |
|---|---|---|---|---|---|---|
| H1 | NLIWLVK | $1.07\times10^{7}$ | $5.35\times10^{6}$ | 0.502 | 0.558 +/- 0.028 | 21.2 +/- 2.8 % |
| | GVTAACPHAGAK | $1.44\times10^{6}$ | $9.59\times10^{5}$ | 0.668 | | |
| | KFKPEIATRPK | $1.18\times10^{6}$ | $6.22\times10^{5}$ | 0.529 | | |
| | FKPEIATRPK | $2.46\times10^{6}$ | $1.32\times10^{6}$ | 0.536 | | |
| H3 | KWDLFVERSK +WDLFVERSK[c] | $5.88\times10^{6}$ | $4.87\times10^{6}$ | 0.827 | 0.720 +/- 0.030 | 27.3 +/- 3.0 % |
| | SSIMRSDAPIGK[d] | $1.05\times10^{7}$ | $7.45\times10^{6}$ | 0.707 | | |
| | LATGMRNVPEK[d] | $6.24\times10^{6}$ | $3.99\times10^{6}$ | 0.638 | | |
| | EFSEVEGRIQDLEK[e] | $8.62\times10^{6}$ | $6.11\times10^{6}$ | 0.709 | | |
| B/Yam | IRLSTQNVIDAEK | $5.44\times10^{6}$ | $3.77\times10^{6}$ | 0.693 | 0.632 +/- 0.012 | 24.0 +/- 1.6 % |
| | SLYGDSNPQK | $5.01\times10^{6}$ | $3.11\times10^{6}$ | 0.621 | | |
| | SYFANLK | $6.73\times10^{6}$ | $4.00\times10^{6}$ | 0.595 | | |
| | IRQLPNLLRGYEK | $1.30\times10^{7}$ | $8.07\times10^{6}$ | 0.620 | | |
| B/Vic | GSLPLIGEADCLHEK | $7.38\times10^{6}$ | $5.19\times10^{6}$ | 0.703 | 0.727 +/- 0.013 | 27.6 +/- 1.3 % |
| | LYGDSKPQK | $4.92\times10^{5}$ | $3.60\times10^{5}$ | 0.731 | | |
| | TGTITYQRGILLPQK | $6.72\times10^{6}$ | $4.70\times10^{6}$ | 0.700 | | |
| | VWCASGRSK | $1.99\times10^{6}$ | $1.54\times10^{6}$ | 0.773 | | |

[a]Only abundant peptides that were unique to each specific HA were included.
[b]Normalized to the total sum of the average relative ratios.
[c]A high degree of missed cleavage was observed for this peptide and the sum of the two peptides was used for the calculations.
[d]The oxidized methionine forms were also included in these calculations.
[e]The pyro-glutamic acid form of the peptide was also included in these calculations.

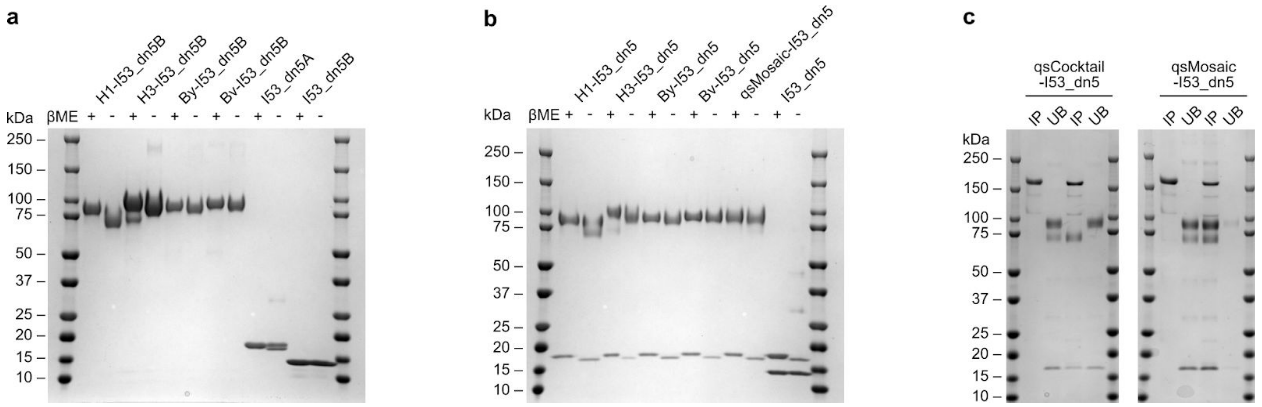


**Supplementary Fig. 1 | Uncropped SDS-PAGE of H1-I53_dn5B fusions, purified nanoparticles and immunoprecipitations. a,** Uncropped SDS-PAGE gel from Extended Data Fig. 1b. **b,** Uncropped SDS-PAGE gel from Extended Data Fig. 1e. **c,** Uncropped SDS-PAGE gel of qsCocktail-I53_dn5 and qsMosaic-I53_dn5 immunoprecipitations from Fig. 1f. IP, immunoprecipitated; UB, unbound.
